# An RNA relay to signal mitochondrial defects across tissues at the cost of healthy aging

**DOI:** 10.64898/2026.11.26.769611

**Authors:** Carmen M. Ramos, Juan I. Bravo, Marisa Gaglio, Tripti Nair, Nicole L. Stuhr, Chia-An Yen, Claire Chung, Dilawar Mir, Marc Vermulst, Sean P. Curran

## Abstract

Mitochondrial dysfunction is a hallmark of aging, yet how stressed mitochondria communicate across tissues to coordinate organismal responses remains poorly understood. Here, we identify an RNA-mediated signaling relay that links mitochondrial dysfunction to tissue-specific and cell non-autonomous aging trajectories. Loss of the mitochondrial 1-pyrroline-5-carboxylate catabolism enzyme ALH-6 leads to adenosine deaminase acting on RNA (ADR-2) activity within the pharyngeal muscle that reduces healthspan and lifespan. RNA editing is further enhanced with age, particularly among transcripts regulated by the mitochondrial RNA polymerase, implicating mitochondrial RNA metabolism as a source of stress-associated RNA signals. We further show that DRH-1, the *Caenorhabditis elegans* orthologue of the RIG-I-like RNA sensors, functions outside the pharyngeal muscle to mediate organism-wide effects on healthspan and longevity, demonstrating that RNA-dependent mitochondrial stress signaling extends beyond the tissue of origin. Consistent with this model, the double-stranded RNA transporter SID-1 facilitates signaling from the pharynx to peripheral tissues. Together, these findings establish RNA editing and inter-tissue RNA-based signaling as integral components of a mitochondrial surveillance pathway that coordinates organismal adaptation to mitochondrial dysfunction. Our work identifies edited RNA as both the intracellular effector and molecular messenger of mitochondrial stress, revealing a mechanism where localized mitochondrial dysfunction drives systemic aging phenotypes.

## INTRODUCTION

Mitochondria are multifunctional organelles that coordinate cellular metabolism, bioenergetics, and stress signaling. Beyond their canonical role in ATP production through oxidative phosphorylation, mitochondria integrate metabolic and environmental cues to regulate redox homeostasis, apoptosis, and adaptive stress responses^1–3^. Consequently, mitochondrial dysfunction, characterized by impaired oxidative metabolism and elevated reactive oxygen species (ROS), is a defining feature of aging and contributes to the pathogenesis of numerous age-associated disorders, including neurodegenerative disease and metabolic dysfunction^4–6^. Despite extensive evidence linking mitochondrial decline to aging, the molecular mechanisms by which mitochondrial metabolic perturbations engage cellular signaling pathways to influence organismal healthspan remain incompletely understood.

One metabolic pathway increasingly implicated in mitochondrial quality control is amino acid catabolism^7^. Within mitochondria, proline is oxidized to 1-pyrroline-5-carboxylate (P5C) by proline dehydrogenase (PRDH-1), whereas arginine is converted to ornithine by arginase (ARGN-1) before ornithine aminotransferase (OATR-1) generates P5C. Mitochondrial 1-pyrroline-5-carboxylate dehydrogenase (ALH-6/P5CDH) catalyzes the final step of arginine, ornithine, and proline degradation by converting P5C to glutamate^8,9^. In mammals, the ALH-6 orthologue, ALDH4A1, has been associated with hyperprolinemia, atherosclerosis, and schizophrenia^10,11^, and we previously demonstrated that genetic variation in ALDH4A1 is a strong predictor of age-associated muscle decline in humans^12^. In *Caenorhabditis elegans*, loss of *alh-6* selectively activates the oxidative stress reporter *gst-4p::GFP* within body-wall muscle, revealing a tissue-specific mitochondrial stress response^8,9,12^. These animals exhibit premature aging characterized by metabolic dysfunction and reduced lifespan^8^. Mechanistically, loss of ALH-6 leads to mitochondrial accumulation of P5C, promoting ROS production and activation of the conserved redox-responsive transcription factor SKN-1/NRF2, thereby driving progressive deterioration in healthspan and lifespan^8,9,12^.

In parallel with their metabolic functions, mitochondria have emerged as central regulators of innate antiviral immunity^13^. Viral infection frequently perturbs mitochondrial homeostasis, and host cells have evolved surveillance mechanisms that detect mitochondrial dysfunction as a trigger for immune activation^13^. In mammals, mitochondrial stress can stimulate antiviral responses through the RIG-I-like receptor (RLR) pathway, in which the RNA helicase RIG-I recognizes viral double-stranded RNA (dsRNA) and initiates interferon signaling^14^. Although *C. elegans* lacks canonical interferon pathways, it possesses the RIG-I-like helicase DRH-1, which detects cytosolic dsRNA and coordinates antiviral defense^15^. DRH-1 is required for transcriptional immune responses during viral infection and contributes to activation of the intracellular pathogen response (IPR) following infection with intracellular pathogens^16^. More recently, we identified an additional role for DRH-1 in the intestine, where it suppresses age-associated pathology driven by constitutive SKN-1 activation^17^, suggesting that RNA surveillance pathways contribute more broadly to mitochondrial stress adaptation during aging.

To prevent inappropriate activation of antiviral signaling, endogenous dsRNAs are extensively modified by adenosine deaminases acting on RNA (ADARs), which catalyze adenosine-to-inosine (A-to-I) editing within double-stranded RNA^18,19^. This editing destabilizes dsRNA duplexes, preventing recognition by RIG-I and related RNA sensors while simultaneously influencing coding potential, alternative splicing, microRNA targeting, and transcript stability^20^. In *C. elegans*, loss of ADAR activity (*adr-1; adr-2* double mutants) results in hyperactive RNA interference, enhanced transgene silencing, and shortened lifespan, underscoring the importance of RNA editing in maintaining transcriptome integrity and organismal health^21^. Moreover, ADAR activity is dynamically regulated by diverse cellular stressors, including hypoxia, heat shock, and mitochondrial dysfunction^22^, linking RNA editing directly to stress adaptation. Consistent with this, dysregulation of RIG-I signaling and ADAR1/2 activity contributes to chronic inflammation and immune dysfunction in mammals^13,23^, highlighting an evolutionarily conserved interface between mitochondrial homeostasis, RNA metabolism, and innate immunity.

Here, we investigated whether disruption of mitochondrial P5C metabolism engages dsRNA surveillance pathways to regulate aging. Building on established roles for ADR-2 and DRH-1 in antiviral defense and the emerging connection between mitochondrial dysfunction and innate immunity, we tested whether mitochondrial metabolic stress alters RNA editing to influence organismal healthspan. We find that mitochondrial dysfunction increases ADR-2-dependent RNA editing and identify a previously unrecognized role for RNA editing in modulating stress responses during aging. Tissue-specific restoration of *alh-6* defined the cellular requirements for mitochondrial amino acid catabolism in healthy aging, whereas tissue-specific expression of *adr-2* revealed that RNA editing within the pharynx unexpectedly promotes lifespan shortening following loss of *alh-6*. These findings support a model in which disruption of mitochondrial P5C catabolism induces a chronic antiviral-like state that is normally restrained under physiological conditions, consistent with previous observations that DRH-1 colocalizes with dsRNA during viral infection^24^. Together, our results identify an unexpected signaling axis linking mitochondrial metabolism, RNA editing, and antiviral surveillance, revealing an integrated stress-response network that coordinates mitochondrial quality control with organismal aging.

## RESULTS

### Cell non-autonomous responses to impaired clearance of mitochondrial P5C

The mitochondrial aldehyde dehydrogenase ALH-6 catalyzes the final step of proline and arginine catabolism, converting the reactive intermediate 1-pyrroline-5-carboxylate (P5C) into glutamate, a metabolic reaction essential for cellular homeostasis^8^. Loss-of-function mutations in *alh-6*, the *Caenorhabditis elegans* ortholog of human *ALDH4A1*, disrupts P5C clearance and leads to reduced lifespan and healthspan^9,12,25–28^ (**Figure 1A**). Notably, *alh-6* loss-of-function (*alh-6lf*) mutants were originally identified through the age-dependent activation of the oxidative stress reporter *gst-4p::GFP*, which becomes selectively induced in body wall muscle (BWM) after the onset of adulthood^8^. Because ALH-6 is expressed broadly across multiple tissues^8,29^, we asked whether disrupted mitochondrial P5C catabolism functions locally or instead elicits tissue-to-tissue communication that influences organismal physiology.

**FIGURE 1.**
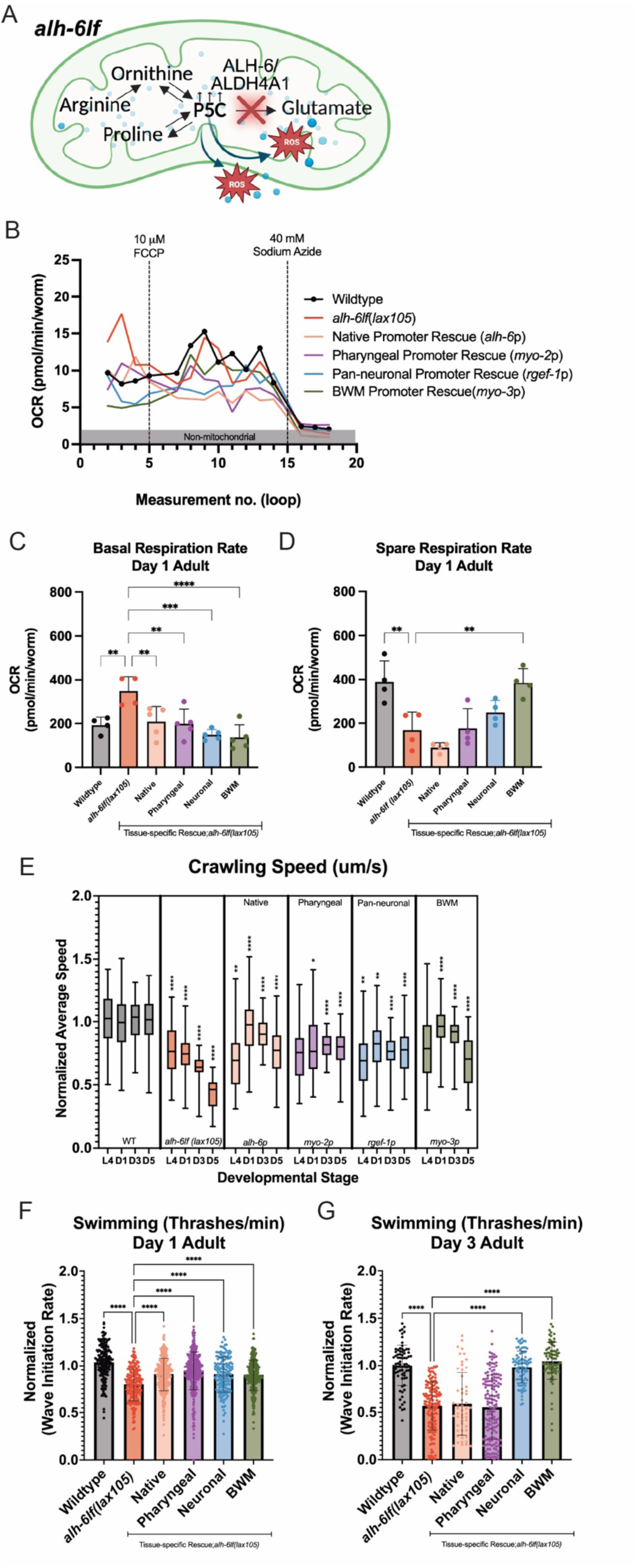
Cell non-autonomous surveillance of mitochondrial P5C influences age-related muscle decline. **(A)** Loss of *alh-6* activates the SKN-1/NRF2 stress response, accelerating mitochondrial and muscle dysfunction in *C. elegans* due to accumulation of the toxic metabolic intermediate, pyrroline-5-carboxylate (P5C). To assess the functionality of the tissue-specific *alh-6* rescues in maintaining mitochondrial health **(B-D)** and muscle mobility **(E-G)**, animals were analyzed throughout adulthood [larval stage 4 (L4) through Day 5 (D5)]. **(B)** Whole-animal oxygen consumption rate (OCR) was measured as a proxy for mitochondrial respiratory activity. Day 1 *alh-6lf* animals showed **(C)** elevated basal respiration rates (**D)** and reduced spare respiration capacity, indicative of overworked and energetically stressed mitochondria. **(E)** *alh-6lf* animals displayed early-onset muscle dysfunction relative to WT; tissue-specific expression of *alh-6* in the pharynx, neurons, or BWM partially rescued age-dependent movement defects. **(F-G)** Thrashing speeds declined with age in *alh-6lf* animals but were significantly restored to WT levels in neuronal and BWM rescues. Together, these findings show a cell non-autonomous contribution to mitochondrial and muscle homeostasis. All raw data counts for OCR can be found in Table S1, and crawling and swimming speed data counts can be found in Table S2. For OCR analyses, n=10-20 animals per strain, N=4. For movement analyses, n=50-100 animals per strain, N=3. Statistical comparisons were performed using one-way ANOVA. *p<0.05, **p<0.005, **** p<0.0001.

To address this question, we generated a panel of transgenic *alh-6lf* strains expressing wild-type *alh-6* (*alh-6wt*) under tissue-specific promoters corresponding to the principal sites of endogenous *<u>alh-6</u>* expression (Figure S1)^9^. These included the pharyngeal muscle (*myo-2p*), pan-neuronal (*rgef-1p*), and body wall muscle (*myo-3p*) promoters. As a physiological control, we also restored *alh-6wt* expression using the endogenous *alh-6* promoter (*alh-6p*).

We first examined whether tissue-specific restoration of *alh-6wt* was sufficient to suppress the age-dependent activation of *gst-4p::GFP* in *alh-6lf* animals. Expression of *alh-6wt* from an endogenous promoter sequence completely restored normal reporter activity (Figure S1A–G). Unexpectedly, expression of *alh-6wt* in each individual tissue—including the pharyngeal muscle and neurons—was also sufficient to suppress *gst-4p::GFP* activation within the body wall muscle (Figure S1A–G). Although body wall muscle-specific rescue (*myo-3p*) is consistent with a cell-autonomous requirement for ALH-6, suppression mediated by pharyngeal or neuronal expression demonstrates that mitochondrial P5C metabolism initiates signaling between tissues that regulates stress responses at distal sites. These findings identify a robust cell non-autonomous pathway through which mitochondrial metabolic status is communicated across tissues to coordinate organismal stress responses.

### Tissue-specific mitochondrial P5C catabolism differentially impacts muscle health and mitochondrial function

Mitochondrial dysfunction during aging is accompanied by progressive alterations in mitochondrial morphology and network organization that reflect declining organelle homeostasis^30^. Because disruption of mitochondrial architecture contributes directly to age-associated muscle dysfunction and disease susceptibility^31^, we first examined age-dependent changes in mitochondrial morphology in wild-type and *alh-6lf* animals expressing a mitochondrially targeted GFP reporter^32^. Mitochondrial networks in both genotypes exhibited the expected continuum of morphologies, ranging from discrete organelles to interconnected tubular networks; however, the temporal progression of these structural changes differed markedly between genotypes (Figure S2A–P). By day 5 of adulthood, *alh-6lf* animals displayed increased mitochondrial interconnectivity within body wall muscle, whereas by day 9, mitochondria adopted a fragmented, spherical morphology characteristic of mitochondrial dysfunction^33^.

To determine how tissue-specific mitochondrial P5C catabolism influences mitochondrial physiology, we quantified oxygen consumption rates (OCR) in the tissue-specific *alh-6* rescue strains. *alh-6lf* mutants exhibited significantly elevated basal respiration relative to wild-type animals, and restoration of *alh-6wt* expression in any of the examined tissues normalized basal respiratory activity (**Figure 1B,C** and Table S1). In contrast, *alh-6lf* mutants displayed a marked reduction in spare respiratory capacity, indicative of diminished metabolic reserve, and this defect was rescued exclusively by body wall muscle-specific expression of *alh-6lf* (**Figure 1B,D**, Figure S2Q and Table S1). These findings indicate that basal mitochondrial metabolism is regulated through both cell-autonomous and cell non-autonomous mechanisms, whereas maintenance of maximal respiratory capacity requires ALH-6 activity within muscle.

We next asked whether these tissue-specific effects on mitochondrial physiology translate into functional improvements in muscle performance. Crawling and swimming behaviors provide sensitive, age-dependent measures of muscle function in *C. elegans*^34,35^. Consistent with previous studies^8,12^, *alh-6lf* mutants exhibited a premature decline in both crawling and swimming performance (**Figure 1E–G** and Table S2). Restoration of *alh-6wt* under the endogenous promoter improved crawling speed at the onset of adulthood but failed to prevent its progressive decline with age, a phenotype similarly observed following body wall muscle-specific rescue (**Figure 1E**). In contrast, expression of *alh-6wt* in either the pharyngeal muscle or neurons preserved crawling performance throughout the five-day analysis (**Figure 1E**). Swimming performance showed a distinct tissue dependence. Although all rescue lines modestly improved swimming speed on day 1 of adulthood (**Figure 1F**), only neuronal and body wall muscle expression restored swimming capacity to wild-type levels by day 3 (**Figure 1G**).

Together, these findings demonstrate that mitochondrial P5C catabolism exerts tissue-specific effects on mitochondrial function and muscle physiology. While restoration of ALH-6 activity in multiple tissues is sufficient to normalize systemic metabolic stress responses, distinct tissues make non-equivalent contributions to the preservation of muscle function during aging, supporting the existence of specialized cell non-autonomous signaling pathways that coordinate organismal responses to mitochondrial dysfunction.

### Mitochondrial P5C catabolism in the pharyngeal muscle is required for normal lifespan

Because *alh-6lf* mutations accelerate organismal aging^8,9,12,28^, we next asked whether tissue-specific restoration of ALH-6 could rescue the shortened lifespan associated with defective mitochondrial P5C catabolism. Expression of *alh-6wt* under its endogenous promoter partially restored longevity by increasing both median and maximal lifespan; however, survival during the first quartile of life remained indistinguishable from that of *alh-6lf* mutants (**Figure 2A** and Table S3), indicating that physiological expression of *alh-6* incompletely reverses the aging phenotype.

**FIGURE 2.**
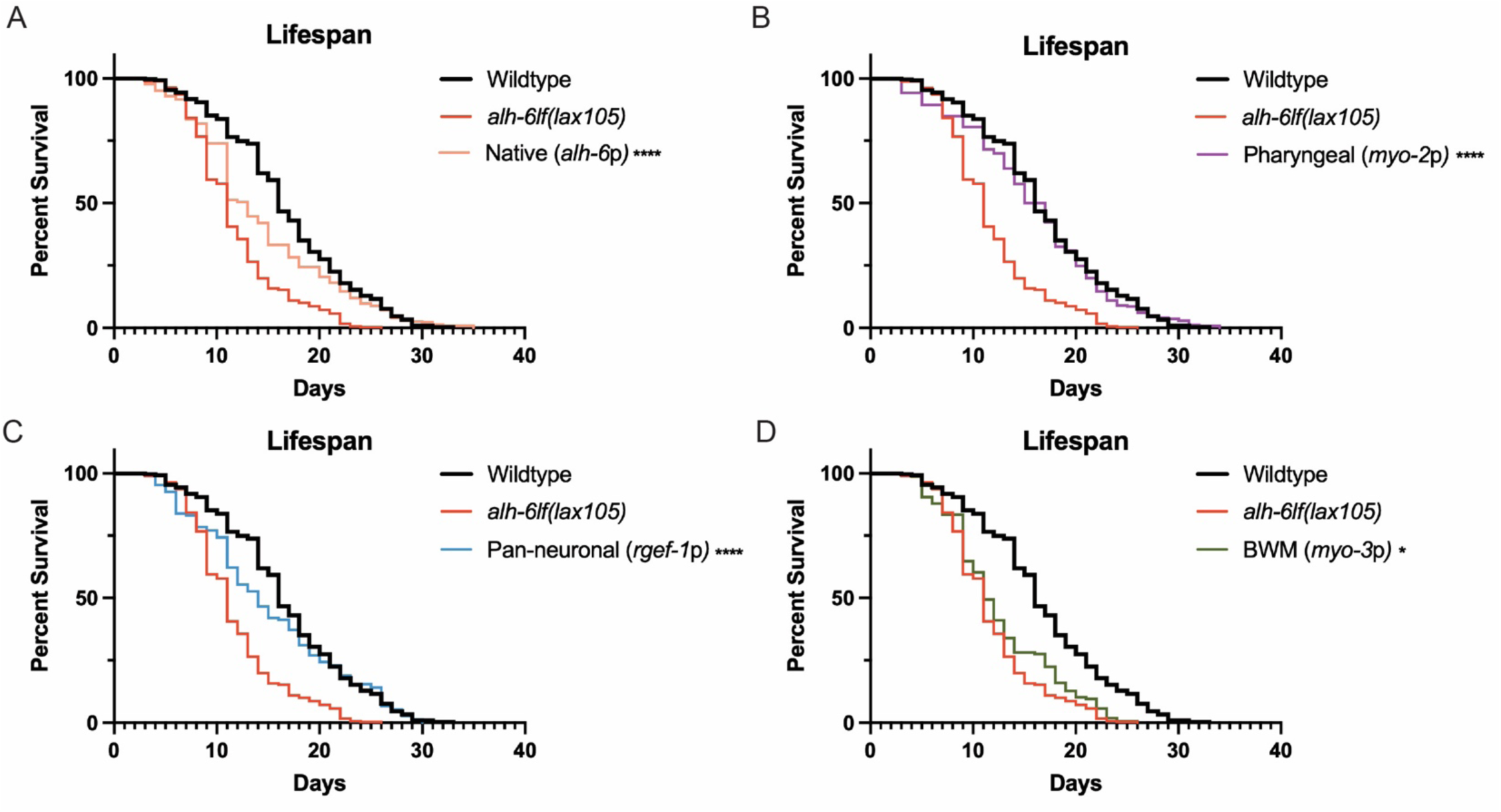
Tissue specific responses to the mitochondrial P5C signaling axis differentially impact longevity. Lifespan assays were performed to assess the impact of tissue-specific *alh-6* rescues on organismal healthspan and survival. Consistent with previous findings, *alh-6lf* animals exhibited a significantly reduced median lifespan of 11 days, compared to WT. **(A)** Native *alh-6* rescue extended median survival to 13 days, with a significant improvement at the third quartile (25% survival), relative to *alh-6lf* animals. **(B)** Pharyngeal and **(C)** pan-neuronal *alh-6* expression significantly improved survival throughout the lifespan, yielding survival curves comparable to that of WT animals. **(D)** BWM-specific rescue modestly extended lifespan, though to a lesser extent than pharyngeal or neuronal expression. Altogether, these findings suggest that cell non-autonomous signals originating from the pharyngeal and neuronal tissues play key roles in coordinating survival. All statistical analyses were performed using Log-rank (Mantel-Cox) test, found in Table S3; tissue-specific rescues were compared to *alh-6lf*. *p<0.05, **** p<0.0001. For lifespan analyses, n=100-150 animals per strain, N=3.

To define the tissue-specific requirements for mitochondrial P5C catabolism, we assessed lifespan following targeted restoration of *alh-6wt*. Remarkably, expression in the pharyngeal muscle (*myo-2p*) significantly extended lifespan, restoring survival toward wild-type levels (**Figure 2B** and Table S3). Pan-neuronal expression (*rgef-1p*) similarly improved longevity (**Figure 2C** and Table S3). In contrast, body wall muscle-specific expression (*myo-3p*) failed to rescue lifespan despite effectively suppressing *gst-4p::GFP* activation (Figure S1) and restoring multiple indices of mitochondrial function and muscle health (**Figure 1**; **Figure 2D** and Table S3).

These findings reveal that correction of local mitochondrial dysfunction within body wall muscle is insufficient to restore organismal longevity. Instead, lifespan is selectively influenced by mitochondrial P5C catabolism in specific tissues, most prominently the pharyngeal muscle. Together with the tissue-specific suppression of stress signaling (Figure S1), these data demonstrate that distinct tissues contribute differentially to healthspan and lifespan and identify the pharyngeal muscle as a critical regulator of organismal aging in response to mitochondrial metabolic dysfunction. The ability of pharyngeal ALH-6 to influence longevity independently of body wall muscle physiology further supports the existence of a cell non-autonomous signaling pathway that communicates mitochondrial metabolic status between tissues to coordinate aging at the organismal level.

### RNA editing mediates the muscle response to defective mitochondrial P5C catabolism

To identify regulators of the mitochondrial stress response elicited by defective P5C catabolism, we performed an unbiased ethyl methanesulfonate (EMS) suppressor screen in *alh-6lf*;*gst-4p::GFP* animals, selecting for mutants that failed to induce the characteristic age-dependent activation of the stress reporter in body wall muscle. This strategy parallels our previous genetic screen, which identified *prdh-1*, encoding the enzyme catalyzing the first step of proline catabolism upstream of ALH-6 (**Figure 1A**)^28^. From this screen, we isolated a strong suppressor allele, *lax229*, which abolished *gst-4p::GFP* activation in the body wall muscle at Day 3 of adulthood (**Figure 3A,B**).

**FIGURE 3.**
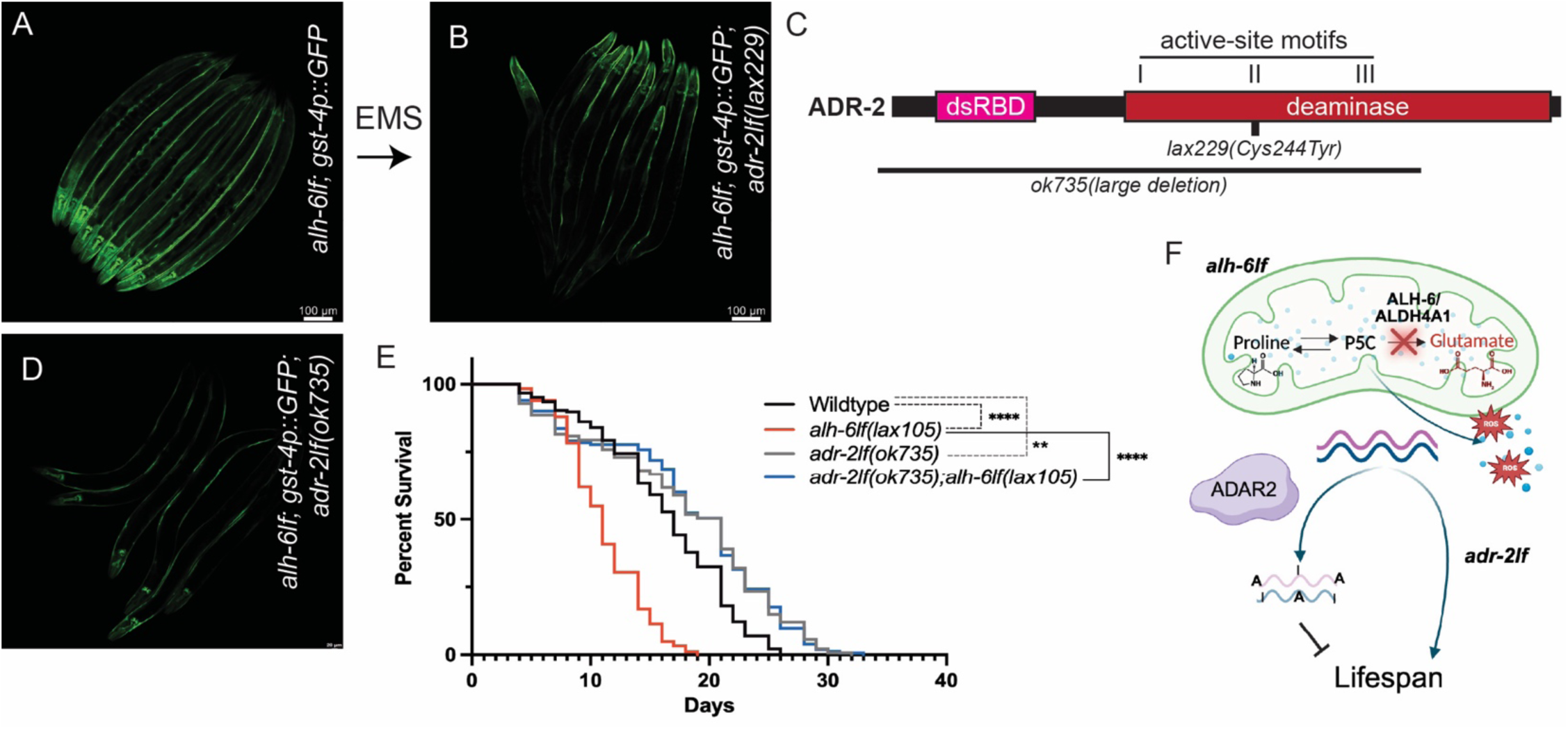
RNA editing drives premature aging in mitochondrial P5C catabolism mutants. (A-B) An EMS mutagenesis screen for suppressors of the adult and muscle-specific activation of the *gst-4p::GFP* reporter identified *adr-2lf(lax229)* as a suppressor of *alh-6lf;gst-4p::GFP* animals. **(C)** Whole-genome sequencing and InDel mapping identifies genetic linkage of *lax229* to the center of chromosome III, revealing a missense mutation converting a conserved cysteine to tyrosine within the catalytic cite of the ADR-2 deaminase domain. **(D)** Using a characterized null allele, *adr-2(ok735),* which contains a large deletion, we confirmed that loss of ADR-2 suppresses the *alh-6lf;gst-4p::GFP* muscle-specific reporter expression. **(E)** Loss of ADR-2 significantly modulates the premature aging phenotype observed in *alh-6lf* mutants, indicating that ADR-2 activity modulates organismal aging downstream to mitochondrial dysfunction (**F**). Statistical analyses were performed using Log-rank (Mantel-Cox) test, found in Table S3. **p<0.005, **** p<0.0001. For lifespan analyses, n=100-150 animals per strain, N=3.

Genetic mapping localized *lax229* to the central region of chromosome III^36–38^, and whole-genome sequencing identified a missense mutation in *adr-2*, which encodes the sole catalytically active adenosine deaminase acting on RNA (ADAR) in *C. elegans*^18,19,39^. The *lax229* mutation substitutes a highly conserved catalytic cysteine (C244) with tyrosine, disrupting the enzymatic domain of ADR-2 (**Figure 3C** and Figure S3A,B). Expression of wild-type *adr-2* (*adr-2wt*) restored *gst-4p::GFP* activation in *alh-6lf*;*lax229;gst-4p::GFP* animals (Figure S3C), confirming that *lax229* is a loss-of-function allele of *adr-2* (*adr-2lf*). Consistent with this conclusion, the independently derived null allele *adr-2(ok735)* also suppressed body wall muscle activation of *gst-4p::GFP* in the *alh-6lf* background (**Figure 3D**). Because *adr-2(ok735)* is a molecular null^21,40^, it was used for all subsequent analyses.

These findings establish that ADR-2-dependent adenosine-to-inosine (A-to-I) RNA editing is required for the BWM stress response triggered by impaired mitochondrial P5C clearance. Since RNA editing has previously been implicated as a negative regulator of longevity in *C. elegans*^21^ we next asked whether loss of ADR-2 activity influences the premature aging phenotype caused by defective mitochondrial P5C catabolism. As reported previously, *adr-2(ok735)* mutants were long-lived relative to wild-type animals (**Figure 3E** and Table S3), but strikingly, deletion of *adr-2* completely suppressed the shortened lifespan of *alh-6lf* mutants,extending lifespan beyond that of wild-type animals (**Figure 3E**and Table S3). The persistence of the longevity phenotype in *alh-6lf*;*adr-2lf* double mutants indicate that ADR-2-dependent RNA editing functions genetically downstream of mitochondrial P5C catabolism to promote age-associated decline.

Together, these genetic analyses identify RNA editing as an essential effector of the mitochondrial stress response initiated by defective P5C metabolism and position ADR-2 as a central mediator linking mitochondrial dysfunction to organismal aging (**Figure 3F**).

### The RNA editing landscape is extensively remodeled by aging and mitochondrial stress

To define how RNA editing changes during aging and in response to defective mitochondrial P5C catabolism, we profiled the editome of wild-type and *alh-6lf* animals at the L4 larval stage and day 7 of adulthood using circle sequencing (circleSeq), a highly accurate approach for detecting adenosine-to-inosine (A-to-I) editing events (**Figure 4A** and Figure S4A)^41–43^. Unlike conventional RNA sequencing methods, circleSeq minimizes false-positive editing calls by requiring editing events to be consistently detected across multiple tandem reads generated from individual circularized RNA molecules^44^. This strategy provides a high-confidence map of age-dependent and genotype-dependent RNA editing events.

**Figure 4.**
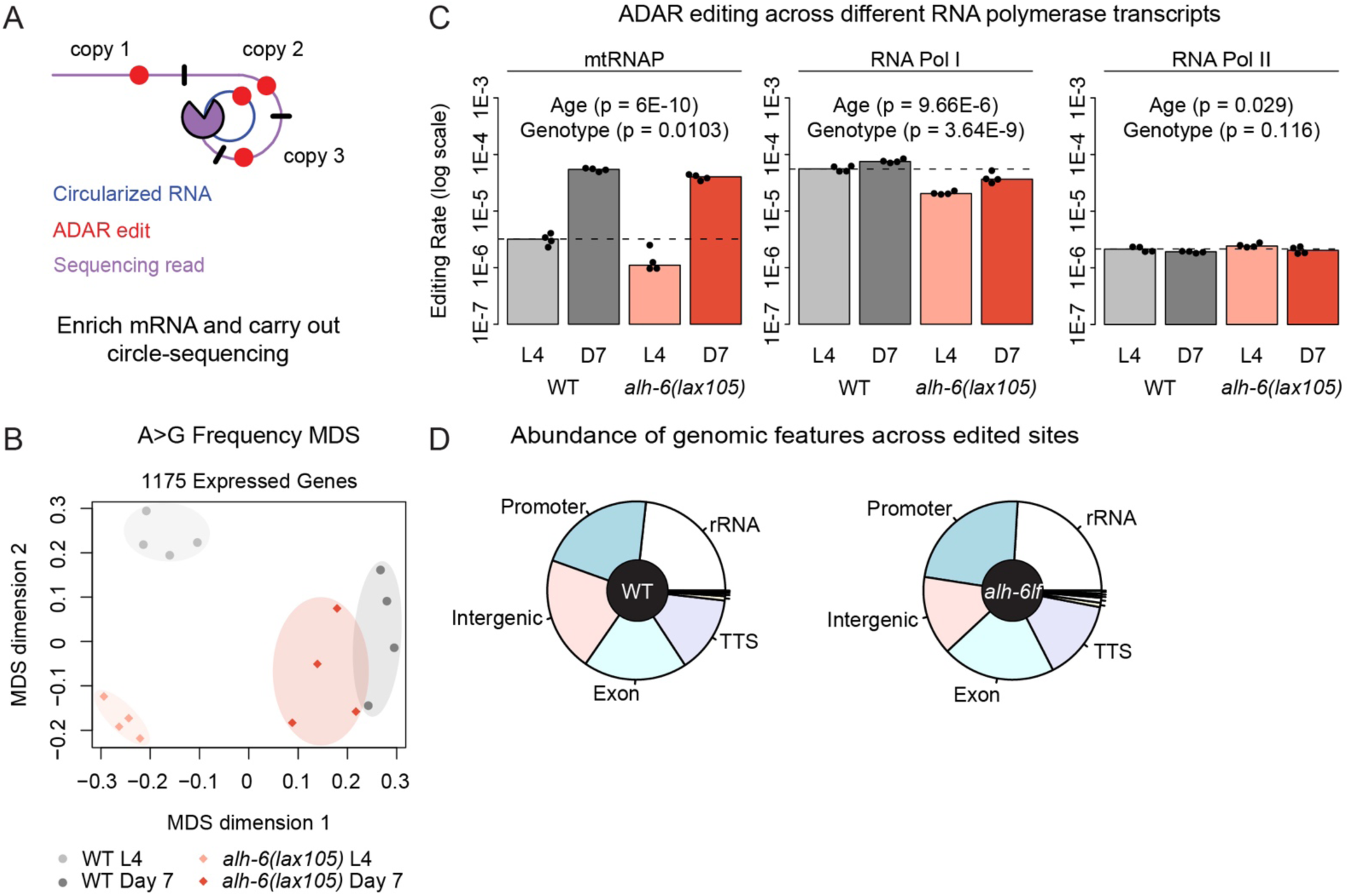
Age and mitochondrial P5C surveillance pathway alter the ADAR editing landscape. **(A)** Diagram illustrating the circle sequencing approach used to characterize ADAR editing events across age and mitochondrial functional status in *C. elegans*. **(B)** Multidimensional scaling (MDS) analysis of gene-level ADAR editing rates in wild-type (WT) and *alh-6lf* worms, with N=4 biological replicates per experimental group. Editing rates for 1175 genes that were detectable in all samples were used for MDS. **(C)** Bar plots for the editing rates across age and genotype groups for transcripts originating from mitochondrial RNA polymerase (mtRNAP), RNA polymerase I (RNA Pol I), and RNA polymerase II (RNA Pol II). Editing in RNA polymerase III (RNA Pol III) transcripts was not detected and therefore no plot is shown for that polymerase. Bar heights were set to the median editing rate within each group. Age and genotype effects were modeled as log10(Mutation Frequency) ∼ Age + Genotype and significance was assessed via rank regression. A p-value < 0.05 was considered significant. **(D)** Pie charts representing the fraction of edited sites belonging to the most abundant genomic and transcriptomic feature annotations. All processed data can be found Table S4.

Across young and aged WT animals, circleSeq identified 2,149 putative A-to-I editing sites distributed among 829 genes (Table S4A). Of these sites, 648 were unique to L4 animals, 1,228 were detected exclusively at day 7, and 273 were shared between both developmental stages. In contrast, only 1,914 edited sites spanning 834 genes were identified in *alh-6lf* animals (Table S4A), including 843 L4-specific sites, 820 day 7-specific sites, and 251 shared sites. While WT animals exhibited a marked increase in the number of detectable ADAR-edited sites with age, *alh-6lf* animals maintained a similar number of detectable ADAR-edited sites with age but shifted to new target sites. These results indicate that aging is accompanied by a broad remodeling of ADAR-mediated editing activity.

To determine whether aging preferentially alters editing within specific transcript classes, we quantified editing frequencies according to the RNA polymerase responsible for transcript synthesis (**Figure 4B** and Table S4B). Detectable editing of mitochondrial RNA polymerase (mtRNAP)-derived transcripts increased significantly with age in both wild-type and *alh-6lf* animals but was attenuated in the *alh-6lf* background. Similarly, RNA polymerase I (RNA Pol I)-derived transcripts, including ribosomal RNAs, exhibited elevated editing frequencies in older animals of both genotypes but lower observable editing in *alh-6lf* mutants compared to wild-type animals. Editing of RNA polymerase II (RNA Pol II)-derived transcripts decreased slightly during aging but trended higher in *alh-6lf*^4645^¾. These findings indicate that both aging and mitochondrial dysfunction selectively reshape ADAR activity across distinct transcript classes. Principal component analysis (PCA) of editing frequencies for 1175 edited genes (Table S4C) further demonstrated that age is the dominant determinant of editome composition. Intriguingly, young WT, young *alh-6lf,* and old *alh-6lf* animals clustered tightly, whereas aged WT samples displayed greater dispersion, consistent with increased heterogeneity in RNA editing during aging (Figure S4B). Furthermore, classical multidimensional scaling (MDS) analysis of 1175 edited genes (Table S4C) revealed a clear separation across both genotype and age in the composition of the editome (**Figure 4C**).

We next examined the genomic distribution of editing events. In both wild-type and *alh-6lf* animals, ADAR editing occurred predominantly within intergenic regions, promoters, exons, ribosomal RNA loci, and transcriptional termination sites (**Figure 4D**). After adjusting for genomic composition, wild-type editing sites were significantly enriched within rRNA loci, intergenic regions, and exons, whereas promoter and intronic sequences were depleted (Table S4D). The *alh-6lf* editome exhibited a similar enrichment profile, with rRNA loci, intergenic regions, exons, and 3’UTRs remaining the most enriched features and promoters and introns remaining depleted (Table S4E). The enrichment of exonic editing likely reflects the high transcript fidelity afforded by circleSeq together with preferential recovery of mature mRNAs during library preparation and then during sequence pseudoalignment, rather than the predominance of intronic editing reported using conventional sequencing approaches.

To determine whether edited transcripts participate in common biological pathways, we performed Gene Ontology enrichment analysis. Edited genes in wild-type animals were significantly enriched for biological processes associated with cellular division, mitochondrial metabolism, and proteostasis, including mitotic cell cycle, oxidative phosphorylation, aerobic respiration, and translation (Figure S4C and Table S4F). Similarly for *alh-6lf* mutants, edited genes remained enriched for mitochondrial metabolic pathways and proteostasis pathways, including translation, cellular respiration, and mitochondrial ATP synthesis coupled to electron transport (Figure S4D and Table S4G).

Collectively, these analyses demonstrate that aging is accompanied by extensive remodeling of the RNA editome and identify mitochondrial metabolic and proteostatic pathways as prominent targets of ADAR-dependent RNA editing. Furthermore, defective mitochondrial P5C catabolism profoundly alters both the magnitude and distribution of RNA editing events, implicating dynamic RNA editing as a component of the adaptive response to mitochondrial dysfunction.

### The pharyngeal muscle is the central site of ADR-2-dependent regulation of organismal aging

The tissue-specific effects of ALH-6 on healthspan and lifespan suggested that RNA editing may similarly act within discrete tissues to mediate the response to defective mitochondrial P5C catabolism. To define the site of ADR-2 action, we restored wild-type *adr-2* (*adr-2wt*) expression in individual tissues of *alh-6lf*;*adr-2lf* animals and assessed the consequences for lifespan.

Remarkably, pharyngeal muscle-specific expression (*myo-2p*) of *adr-2wt* was sufficient to reinstate the shortened lifespan characteristic of *alh-6lf* mutants in *alh-6lf*;*adr-2lf* animals (**Figure 5A** and Table S5), demonstrating that RNA editing within the pharynx is sufficient to drive premature aging in response to impaired mitochondrial P5C catabolism. In contrast, pan-neuronal expression (*rgef-1p*) had no significant effect on lifespan (Figure S5A and Table S5). Body wall muscle-specific rescue (*myo-3p*) produced only a partial phenotype (Figure S5B and Table S5): although median survival shifted toward that of *alh-6lf* animals, and late-life survival more closely resembled that of *alh-6lf*;*adr-2lf* double mutants. These findings identify the pharyngeal muscle as the principal site at which ADR-2-dependent RNA editing promotes organismal aging following disruption of mitochondrial P5C metabolism.

**FIGURE 5.**
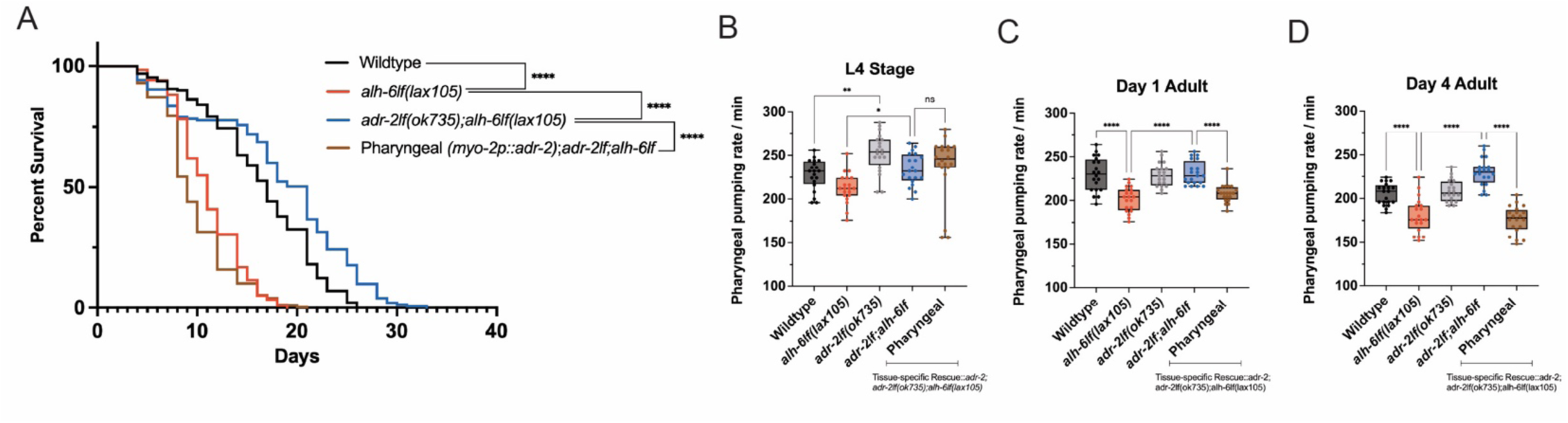
Pharyngeal ADR-2 activity instructs organismal aging. **(A)** Lifespan analysis showing that reintroduction of wildtype *adr-2* in the pharyngeal tissue in *alh-6lf;adr-2lf* animals recapitulate the shortened lifespan phenotype characteristic of *alh-6lf* mutants. Pharyngeal pumping assays performed at **(B)** larval stage 4, **(C)** day 1, **(D)** and day 4 adulthood show that pharyngeal-specific expression of *adr-2wt* in *alh-6lf;adr-2lf* animals recapitulate the age-dependent decline in pumping observed in *alh-6lf* mutants, data counts found in Table S6. Together, these findings demonstrate that pharyngeal RNA editing modulates both lifespan and healthspan in response to disrupted mitochondrial P5C catabolism, acting through a cell non-autonomous mechanism. Lifespan statistical analyses were performed using Log-rank (Mantel-Cox) test, found in Table S5. **** p<0.0001. Pharyngeal pumping statistical comparisons were performed using one-way ANOVA. *p<0.05, **p<0.005, **** p<0.0001. For lifespan analyses, n=100-150 animals per strain, N=3.

Because pharyngeal ADR-2 activity exerted a dominant influence on lifespan, we next asked whether it also regulates age-dependent decline in pharyngeal muscle function. Pharyngeal pumping progressively declined with age in *alh-6lf* animals, as reported previously, whereas loss of *adr-2* substantially preserved pumping activity (**Figure 5B–D** and Table S6). Re-expression of *adr-2wt* specifically in the pharyngeal muscle restored the age-dependent decline in pumping, phenocopying *alh-6lf* animals (**Figure 5B–D** and Table S6). By comparison, neuronal expression of *adr-2* partially improved pharyngeal pumping throughout adulthood, whereas body wall muscle expression failed to reverse the protective effect of *adr-2* loss (Figure S5C,D).

Together, these findings demonstrate that ADR-2-dependent RNA editing functions predominantly within the pharyngeal muscle to couple mitochondrial metabolic dysfunction to organismal aging. The concordant effects on lifespan and pharyngeal muscle function further establish the pharynx as a signaling center that converts local mitochondrial stress into systemic, cell non-autonomous programs regulating healthspan and longevity.

### RIG-I/DRH-1 mediates systemic responses to mitochondrial P5C catabolism across tissues

In mammals, ADAR enzymes regulate innate immune signaling in part by editing endogenous double-stranded RNA (dsRNA) to prevent recognition by RIG-I–like receptors. In *C. elegans*, the RIG-I homolog DRH-1 has been implicated in dsRNA sensing and innate immune responses. We therefore asked whether DRH-1 participates in the systemic response to impaired mitochondrial P5C catabolism.

To test this, we examined whether a gain-of-function allele of *drh-1* (*drh-1gf*, *lax257*), previously identified as a suppressor of constitutive SKN-1 activation^37^, modifies stress signaling in *alh-6lf* animals. Strikingly, *alh-6lf*;*drh-1gf* mutants exhibited a marked reduction in *gst-4p::GFP* activation in body wall muscle (**Figure 6A**), indicating suppression of the stress response. Consistent with this observation, *drh-1gf* also significantly improved survival in *alh-6lf* animals, restoring lifespan to levels comparable to both wild-type and *alh-6lf*;*adr-2lf* mutants (**Figure 6B**). These results position DRH-1 as a second RNA-responsive key regulator of organismal responses to mitochondrial P5C dysfunction.

**FIGURE 6.**
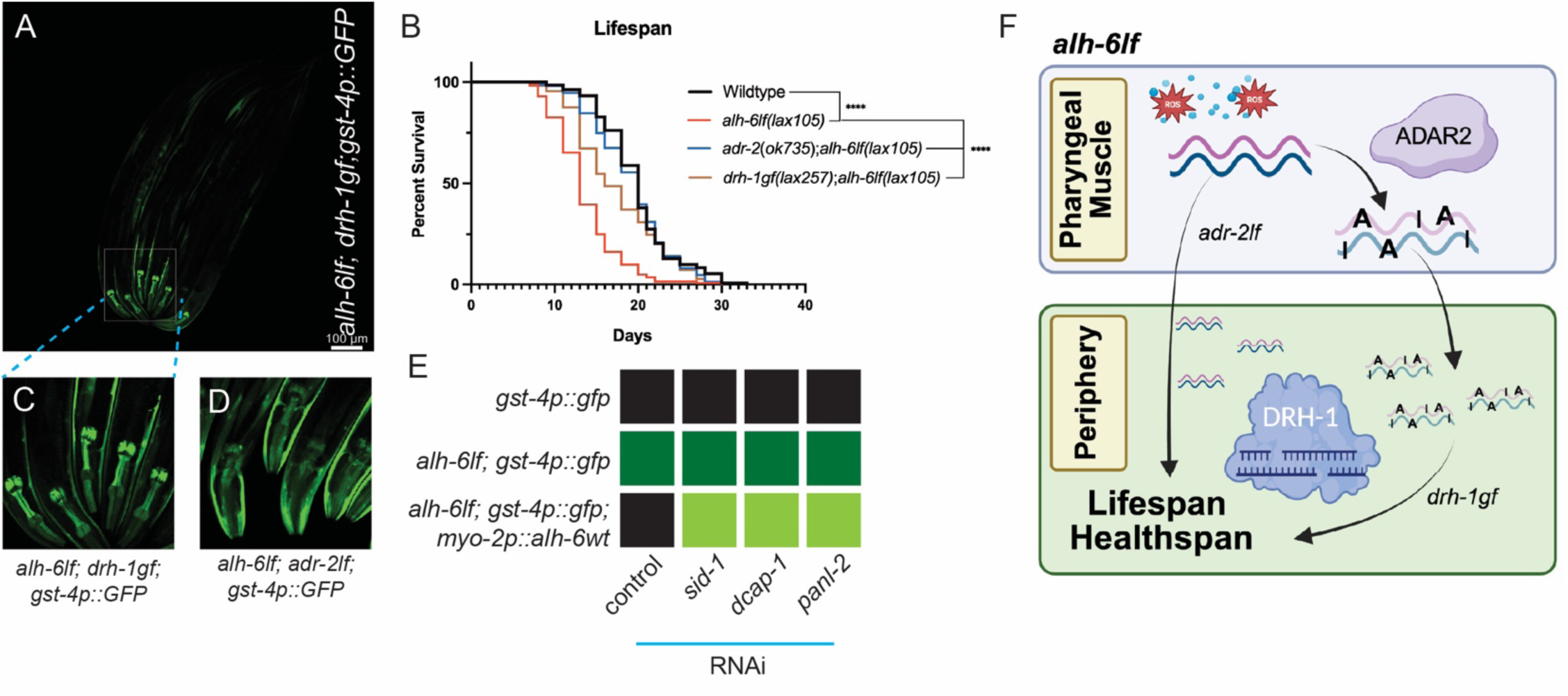
A dsRNA signaling network surveilles systemic stress and longevity responses to mitochondrial dysfunction. **(A)** Constitutive activation of DRH-1 via the gain of function allele *drh-1gf(lax257)* suppresses the *gst-4p::GFP* reporter in the BWM of *alh-6lf* animals. **(B)** The *drh-1gf;alh-6lf* double mutant exhibits a significant lifespan extension, suggesting that enhanced DRH-1 signaling mitigates the mitochondrial stress-induced longevity defect. Suppression patterns of ADR-2 and DRH-1 differ: **(C)** *alh-6lf;drh-1gf* mutants retain strong pharyngeal *gst-4p::GFP* activation, whereas **(D)** *alh-6lf;adr-2lf* animals do not. These results indicate that DRH-1 acts non-autonomously outside the pharynx, while ADR-2 acts cell-autonomously within pharyngeal tissue. **(E)** Heatmap of targeted RNAi screen identified RNA metabolism genes *dcap-1, panl-2,* and the dsRNA transporter *sid-1* as mediators required for distal activation of the *gst-2p::GFP* reporter in *alh-6lf* animals expressing pharyngeal *alh-6wt*. Knockdown of these factors restored *gst-4p::GFP* activity outside the pharynx. (**F**) Cartoon representation where accumulation of a dsRNA-based signals through a network involving the dsRNA-editing enzyme ADR-2, the RNA helicase DRH-1, and the dsRNA transporter SID-1 to coordinate systemic communication of mitochondrial stress from the pharynx to peripheral tissues, modulating longevity. Lifespan statistical analyses were performed using Log-rank (Mantel-Cox) test, n=50-100 animals per strain, N=3. **p<0.005, **** p<0.0001.

Despite the established role of ADARs and RIG-I–like receptors in shared dsRNA pathways within individual cells^47^, the spatial patterns of stress reporter suppression differed between *adr-2lf* and *drh-1gf* mutants in the context of *alh-6lf*. Specifically, *alh-6lf*;*drh-1gf* animals retained robust *gst-4p::GFP* activation in the pharyngeal muscle (**Figure 6C**), whereas this response was largely absent in *adr-2lf* mutants (**Figure 6D**). These findings suggest that ADR-2 functions cell-autonomously within the pharynx to regulate dsRNA-dependent stress signaling, whereas DRH-1 acts predominantly in extra-pharyngeal tissues. This spatial separation implies the existence of a communication axis linking pharyngeal RNA editing activity to systemic DRH-1–dependent stress responses during mitochondrial dysfunction.

To identify additional components of this RNA-based signaling network, we performed a focused RNAi screen targeting genes involved in RNA processing, turnover, and intercellular RNA transport in the context of pharyngeal rescue of *alh-6wt* (*myo-2p::alh-6wt* in an *alh-6lf* background). RNAi-mediated depletion of the mRNA decapping enzyme *dcap-1*, the PAN deadenylation complex subunit *panl-2*, and the dsRNA transporter *sid-1* each restored *gst-4p::GFP* activation outside the pharynx (**Figure 6E** and Figure S6A–L). Notably, *sid-1* encodes a conserved dsRNA channel required for systemic RNA interference, providing functional evidence that dsRNA species coordinate trans-tissue coordination of this systemic stress response.

Together, these findings define a dsRNA-based communication network linking mitochondrial dysfunction in the pharyngeal muscle to organism-wide stress signaling. In this model, ADR-2-dependent RNA editing within the pharynx shapes endogenous dsRNA populations, which are subsequently processed and transmitted through SID-1–dependent transport and interpreted by DRH-1–mediated sensing in distal tissues. This inter-tissue RNA signaling axis integrates mitochondrial metabolic state with systemic stress responses and ultimately determines organismal aging and longevity (**Figure 6F**).

## DISCUSSION

Our study reveals that RNA editing can function not simply as a cell-autonomous response to stress, but as an initiating event in inter-tissue communication that shapes organismal aging. Mitochondrial homeostasis is essential for stress resistance and is a major determinant of organismal aging^48^ and we propose that RNA can serve as a molecular currency for communicating mitochondrial state across tissue boundaries and that an immune-like RNA surveillance system determines whether mitochondrial stress is successfully resolved or translated into organismal decline. Here, tissue-specific rescue of mitochondrial *alh-6* reveals distinct and separable contributions of individual tissues to the response to impaired mitochondrial P5C catabolism. Although restoration of *alh-6* in any tissue examined was sufficient to suppress activation of the *gst-4p::GFP* oxidative stress reporter in body wall muscle, only pharyngeal or neuronal expression restored normal lifespan. This uncoupling of stress signaling from longevity demonstrates that mitochondrial dysfunction engages tissue-specific signaling pathways with distinct physiological outputs and identifies cephalic tissues as principal regulators of organismal aging. Given their exceptional energetic demands and central roles in metabolic regulation, the pharynx and nervous system likely function as metabolic sensors that coordinate systemic responses to mitochondrial stress. Notably, despite restoration of mitochondrial homeostasis, locomotor performance was only partially rescued, suggesting that age-associated muscle dysfunction can become uncoupled from mitochondrial amino acid metabolism or may reflect irreversible damage accumulated before rescue.

Our genetic analyses identify the double-stranded RNA regulators *adr-2* and *drh-1* as key modulators of the response to defective mitochondrial P5C catabolism. The suppression of aging phenotypes by *adr-2* loss-of-function suggests that accumulation of ADAR-edited RNAs contributes directly to disease pathology following disruption of mitochondrial amino acid metabolism. Cell-autonomous mechanisms are vastly overrepresented in the study of mitochondrial stress and recently, the phenylalanyl-tRNA synthetase FARS-1/FARSA was demonstrated to balance longevity and immunity by downregulating endogenous mitochondrial double-stranded RNAs in cultured cells^49^. Here, the pharynx emerged as a critical site initiating this systemic RNA relay system. As one of the most metabolically active organs in *C. elegans*, the pharynx integrates feeding behavior, neuromuscular activity, mitochondrial function, and organismal aging^50^. We find that ADR-2 activity confined to the pharynx is sufficient to promote premature aging, whereas neuronal or body wall muscle expression fails to fully restore the phenotype. These findings identify the pharynx as a previously unrecognized signaling center in which mitochondrial dysfunction promotes maladaptive RNA editing, potentially generating edited double-stranded RNAs or aberrant transcripts that initiate systemic stress responses. Beyond their catalytic activity, ADAR proteins also regulate innate immunity through RNA binding and protein–protein interactions independent of deaminase activity^51^. Determining how these enzymatic and non-enzymatic functions cooperate during mitochondrial stress will be an important avenue for future investigation.

Consistent with this model, transcriptome-wide analysis identified putative ADR-2 editing events across multiple RNA classes, with the strongest aging association occurring in transcripts produced by the mitochondrial RNA polymerase. A limitation of the present study is that CircleSeq libraries incompletely recover tRNAs, which represent established ADAR substrates^43,52^. Nevertheless, CircleSeq provides high-confidence discrimination between library preparation and sequencing errors and authentic RNA editing events, making it well suited for identifying A-to-I editing sites. We also detected increased editing within RNA polymerase I-derived ribosomal RNAs in aged animals. Although ADAR-mediated rRNA editing has not previously been reported in *C. elegans*, the extensive secondary structure and double-stranded regions present within rRNAs ^53^ make them plausible editing substrates. Edited rRNAs, together with mitochondrial and RNA polymerase II transcripts, therefore, represent potential signaling molecules generated during mitochondrial dysfunction. The enrichment of mitochondrial RNA polymerase-derived transcripts is particularly intriguing, as it suggests that RNAs originating from the mitochondrial genome may not be fully recognized as self despite the long evolutionary integration of mitochondria into the eukaryotic cell^47^.

In mammals, ADAR1 prevents inappropriate innate immune activation by marking endogenous double-stranded RNAs as self through A-to-I editing, whereas unedited RNAs are recognized as foreign^54^. Our findings establish an analogous connection between ADR-2-dependent RNA editing in the pharynx and systemic aging through the *C. elegans* RIG-I ortholog DRH-1, a central mediator of antiviral RNA interference and innate immune responses^37,40,55^. Enhanced DRH-1 activity in peripheral tissues, rather than within the pharynx itself, proved essential for resolving the response to defective P5C catabolism originating in the pharynx. These observations reveal a cell non-autonomous pathway in which edited or otherwise aberrant RNAs generated in the pharynx in response to mitochondrial stress, activate DRH-1-dependent signaling in distal tissues, providing a mechanism by which mitochondrial status is communicated throughout the organism.

Our RNAi screen further supports this model by identifying multiple RNA regulatory factors required for systemic stress signaling, including *sid-1*, which mediates systemic RNA transport^56^, *dcap-1*, which regulates mRNA decapping and processing-body dynamics^57^, and *panl-2*, which promotes RNA degradation following deadenylation^58^. Although *C. elegans* lack an RNase L ortholog that degrades RNA downstream of RIG-I signaling in mammals^59,60^, our findings raise the possibility that DCAP-1 and PANL-2 fulfill analogous RNA-processing functions during mitochondrial stress, but notably here in a cell non-autonomous manner. The depletion of *panl-2* increased both nuclear and cytoplasmic GFP accumulation in intestinal cells of the pharyngeal rescue strain, that potentially indicate that tissues beyond body wall muscle can respond to RNA-based signals originating from the pharynx. Together, these observations suggest that multiple layers of RNA transport, surveillance, and degradation participate in coordinating organism-wide adaptation to mitochondrial dysfunction and whether that signal is resolved or progresses to organismal pathology.

Collectively, our findings identify ADR-2-dependent RNA editing as a central regulatory node linking mitochondrial dysfunction to organismal aging. More broadly, our findings challenge the prevailing view that mitochondrial dysfunction influences aging primarily through tissue-autonomous mechanisms. Instead, they demonstrate how a discrete mitochondrial defect can be transformed into a cell-non-autonomous RNA signal that propagates between tissues and ultimately alters the trajectory of healthy aging (**Figure 6F**). This work establishes tissue-specific RNA editing as a critical mediator of mitochondrial stress signaling and reveals an unexpected mechanism by which tissue-specific mitochondrial dysfunction is translated into organism-wide aging. More broadly, these findings suggest that dysregulated recognition of endogenous RNA may represent a conserved mechanism coupling mitochondrial dysfunction to chronic inflammatory signaling and age-associated decline.

## ACKNOWLEDGMENTS

We thank J-F. Gout for technical assistance with circleSeq data analysis, S. Keel, N. Phan, and O. Villa for laboratory assistance and R. West and D. Allred for critical reading of the manuscript. This work was funded by the Hevolution Foundation award HF AGE-004 to SPC, T32AG052374 to JIB and NLS, R25AG076400 to MG, and F31AG090069 to CMR. Some strains were provided by the CGC, which is funded by the NIH Office of Research Infrastructure Programs (P40 OD010440). We thank WormBase for database curation and data access. We also thank the USC School of Gerontology Mitochondria and Metabolism Core Facility for Seahorse measurements of OCR, and the USC School of Gerontology Imaging Core that are both funded in part by the Nathan Shock Center of Excellence P30AG068345.

## Author contributions

Conceptualization: SPC; Methodology: MV and SPC; Investigation: CMR, JIB, MG, TN, CC, DM, NLS, C-AY, and SPC; Visualization: CMR, JIB, and SPC; Supervision: SPC; Writing (original draft): CMR and SPC; Writing (reviewing & editing): CMR, JIB, NLS, MV, and SPC.

## Data and materials availability

The raw circle-sequencing fastq files were deposited at the Sequence Read Archive (SRA) under BioProject PRJNA1520457. All other data are available in the main text or the supplementary materials. The project-specific code used to analyze the circle-sequencing data is available on Github at https://github.com/juanb001/2025_C_elegans_ADAR_Editing/.

**Human Ethics, Animal Ethics, and Consent to Participate declarations**: not applicable.

## SUPPLEMENTARY MATERIALS

### Supplementary Tables

**Supplementary Table S1. Mitochondrial metabolic assays.** Oxygen Consumption Rate (OCR) basal and spare respiration data parameters for *alh-6lf* and tissue-specific *alh-6(wt)* rescues.

**Supplementary Table S2. Body wall muscle function data.** Wormlab movement data parameters for crawling and swimming speed.

**Supplementary Table S3. Tissue specific *alh-6* rescue lifespan assays data.** Mantel-Cox test for *alh-6lf*, tissue-specific *alh-6(wt)* rescue strains, and *adr-2lf(ok735)* survival curves.

**Supplementary Table S4. Identification and enrichment analyses of putative ADAR editing sites.** The lists of editing sites, editing rates, genomic/transcriptomic feature enrichments, and functional enrichments identified from the wild-type and *alh-6lf C. elegans* circle-sequencing.

**Supplementary Table S5. *adr-2lf* lifespan assays data.** Mantel-Cox test for tissue-specific *adr-2(wt)* rescue strains.

**Supplementary Table S6. Pharyngeal muscle function data.** Raw data counts for pharyngeal pumping assays.

### Supplementary Figures

**Supplementary Figure S1.**
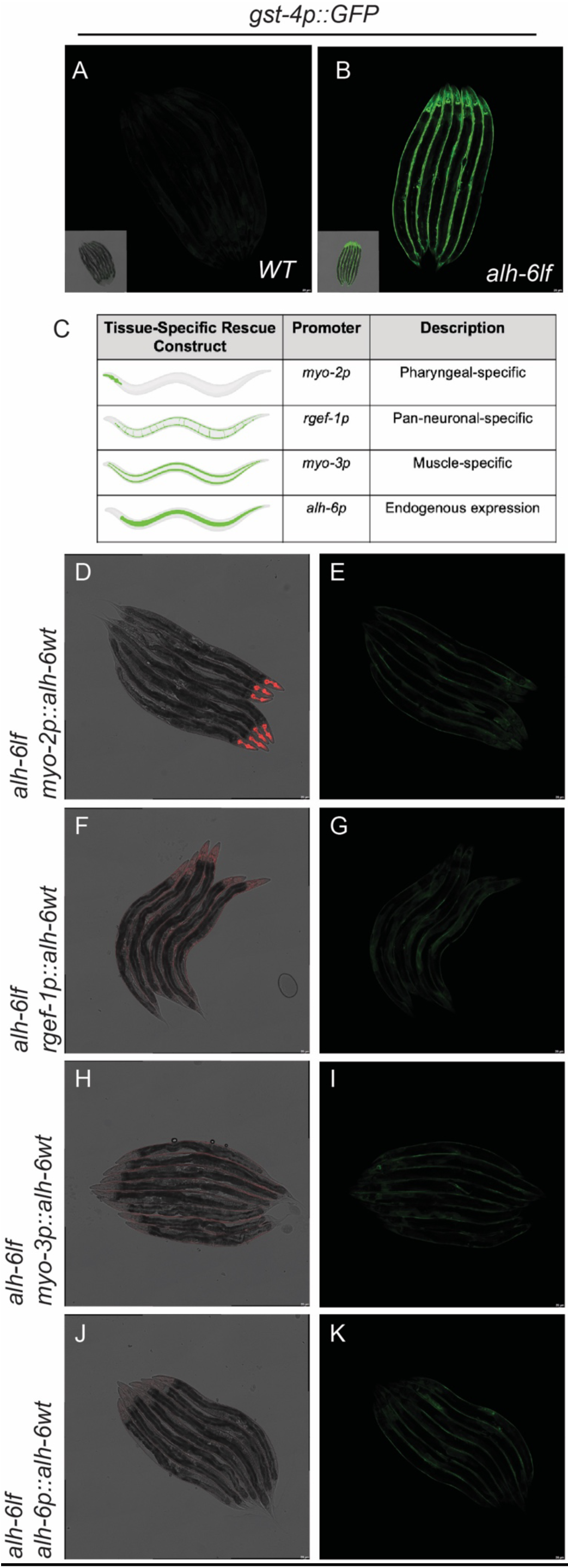
Cell non-autonomous muscle stress responses in mitochondrial P5C catabolism mutants. (**A-B**) BWM-specific activation of the *gst-4p::GFP* reporter in *alh-6lf* mutants is evident by Day 1 of adulthood. (**C**) CRISPR/Cas9 genome editing was used to generate single-copy tissue-specific *alh-6* rescue strains tagged with wrmScarlet to confirm proper transgene integration and expression patterns. <u>(**D-K**)</u> Compared to *alh-6lf* mutants *gst-4p::GFP* expression is suppressed in *alh-6lf* by tissue-specific *alh-6* expression in distal tissues, specifically, (**D-E**) pharyngeal (*myo-2p*), (**F-G**) pan-neuronal (*rgef-1p*), (**H-I)** body wall muscle (*myo-3p*), and (**J-K**) endogenously (*alh-6p*). Together, demonstrating cell non-autonomous regulation of the SKN-1 stress response.

**Supplementary Figure S2.**
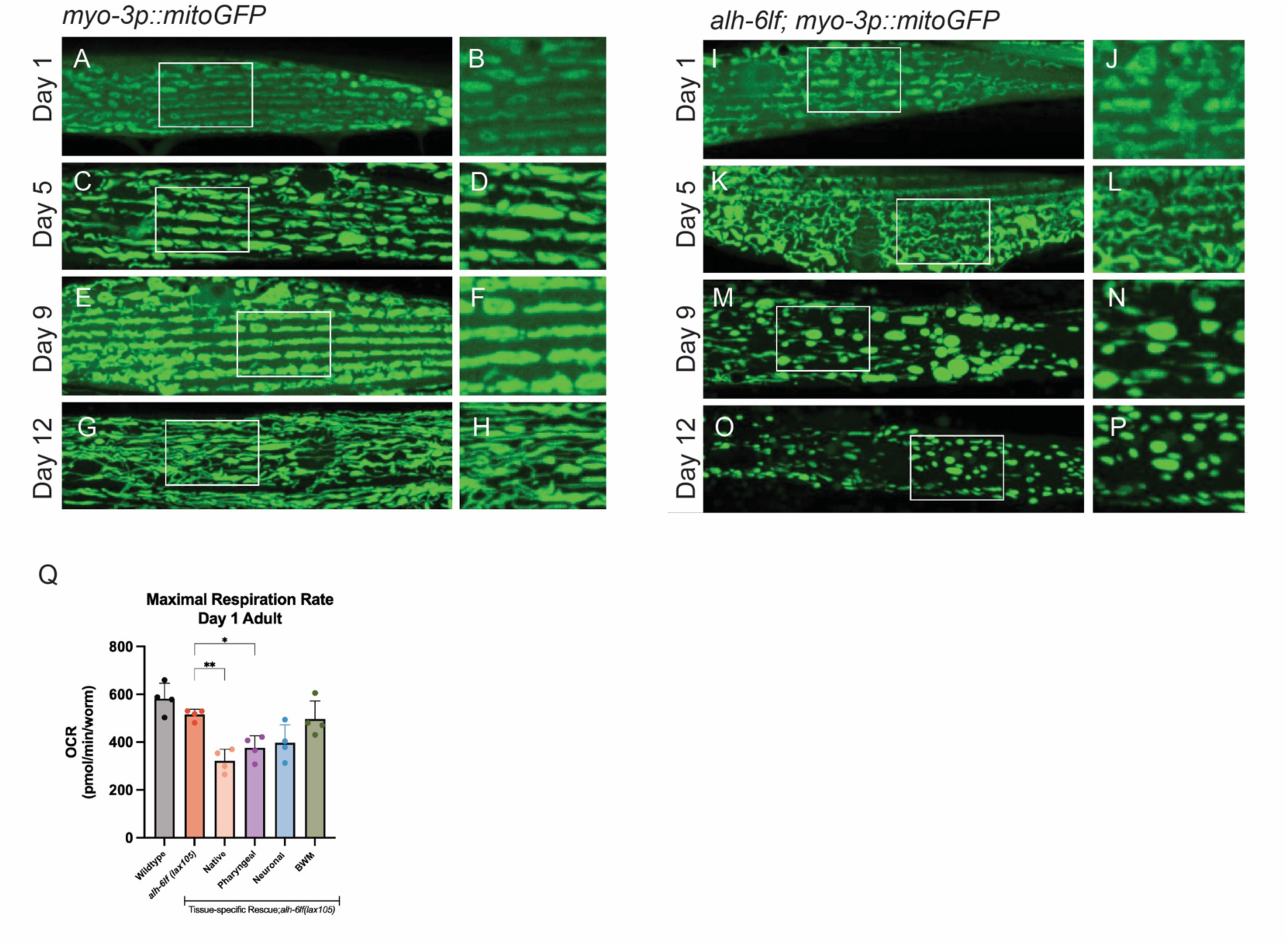
Loss of *alh-6* disrupts mitochondrial morphology and alters respiratory dynamics. **(A-P)** Representative confocal images of mitochondrial morphology in body wall muscle (BWM) of WT and *alh-6lf* animals expressing a mitochondrial-targeted GFP across adulthood. **(Q)** Oxygen consumption rate (OCR) measurements show that maximal respiration rates are maintained, comparable to WT, while expression of *alh-6wt* from a native or pharyngeal promoter results in reduced maximal respiration. These findings show that *alh-6lf* accelerated mitochondrial structure decline with age and modulates respiratory capacity in a tissue-dependent manner.

**Supplementary Figure S3.**
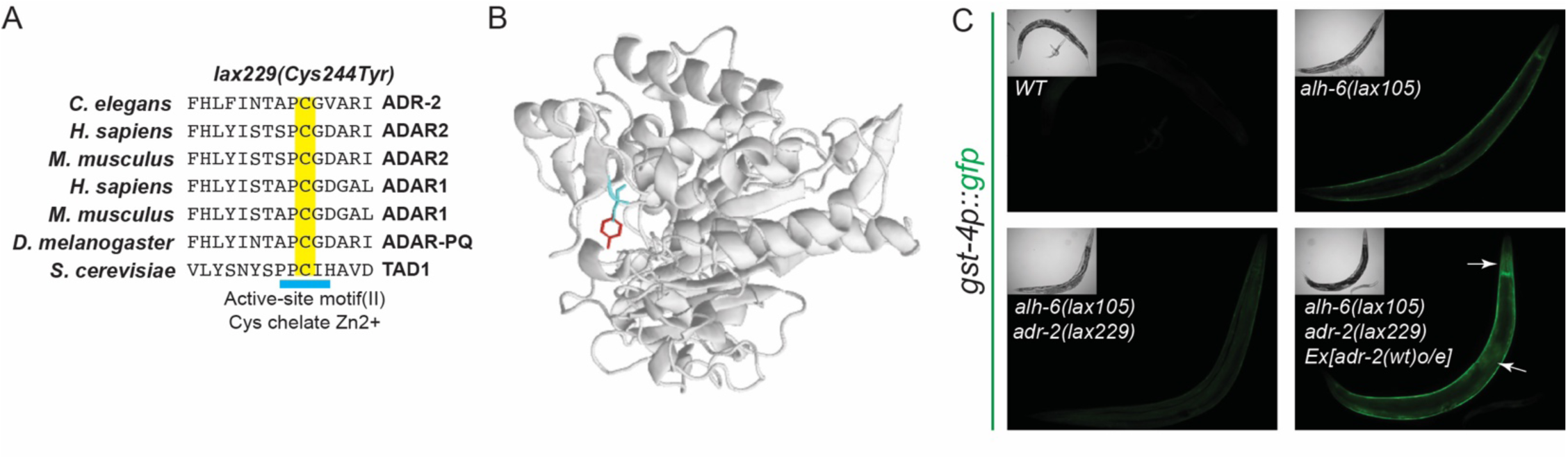
Identification of the *adr-2lf(lax229)* loss-of-function allele. (A-B) Missense mutation in the catalytic domain of *adr-2,* converting a conserved cysteine at position 244 to tyrosine. Sequence alignment shows evolutionary conservation of this cysteine across *C. elegans, H. sapiens, M. musculus, D. melanogaster,* and *S. cerevisiae,* highlighting its functional importance. **(C)** Extrachromosomal expression of wildtype *adr-2 in alh-6lf;adr-2lf(lax229)* mutants restores BWM *gst-4p::GFP* reporter activity, confirming that *lax229* is a loss-of-function allele of *adr-2.* These results validate the identification of *adr-2(lax229)* as a catalytic site mutation responsible for suppression of SKN-1-dependent stress reporter activation in *alh-6lf* animals.

**Supplementary Figure S4.**
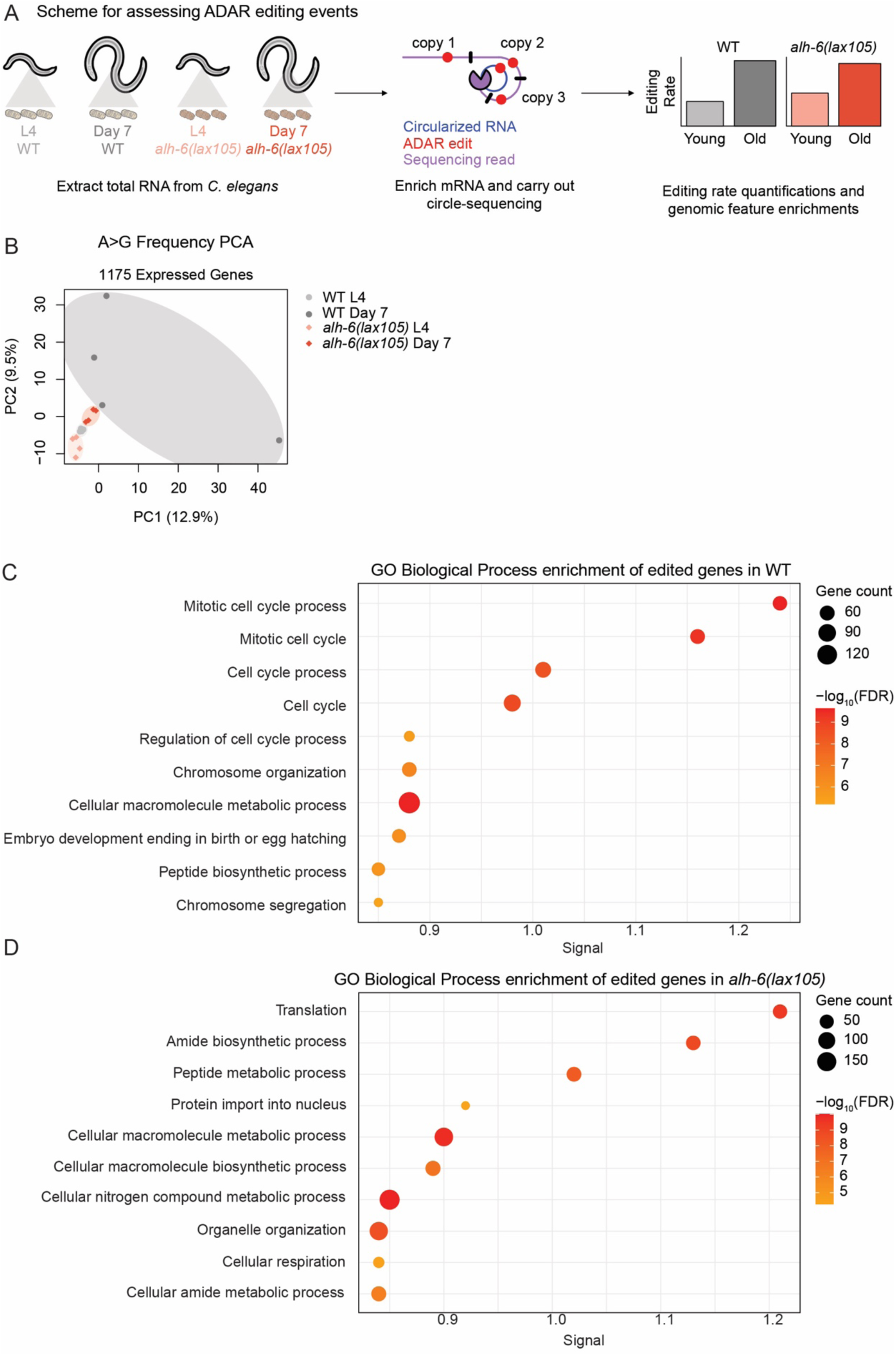
Functional enrichment analysis of ADAR-edited genes. **(A)** Scheme for characterizing ADAR editing events across age and mitochondrial functional status in *C. elegans*. **(B)** Principal component analysis (PCA) of gene-level ADAR editing rates in wild-type (WT) and *alh-6lf* worms, with N=4 biological replicates per experimental group. Editing rates for 1175 genes that were detectable in all samples were used for PCA. Gene Ontology (GO) Biological Process enrichments of ADAR-edited genes identified in the **(C)** wild-type (WT) and **(D)** *alh-6lf* worms. Biological Processes with an FDR < 0.05 were considered significant. FDR: False Discovery Rate. See Table S4 for the complete list of results. Some artwork for panel **(A)** was taken from Bioicons (bioicons.com).

**Supplementary Figure S5.**
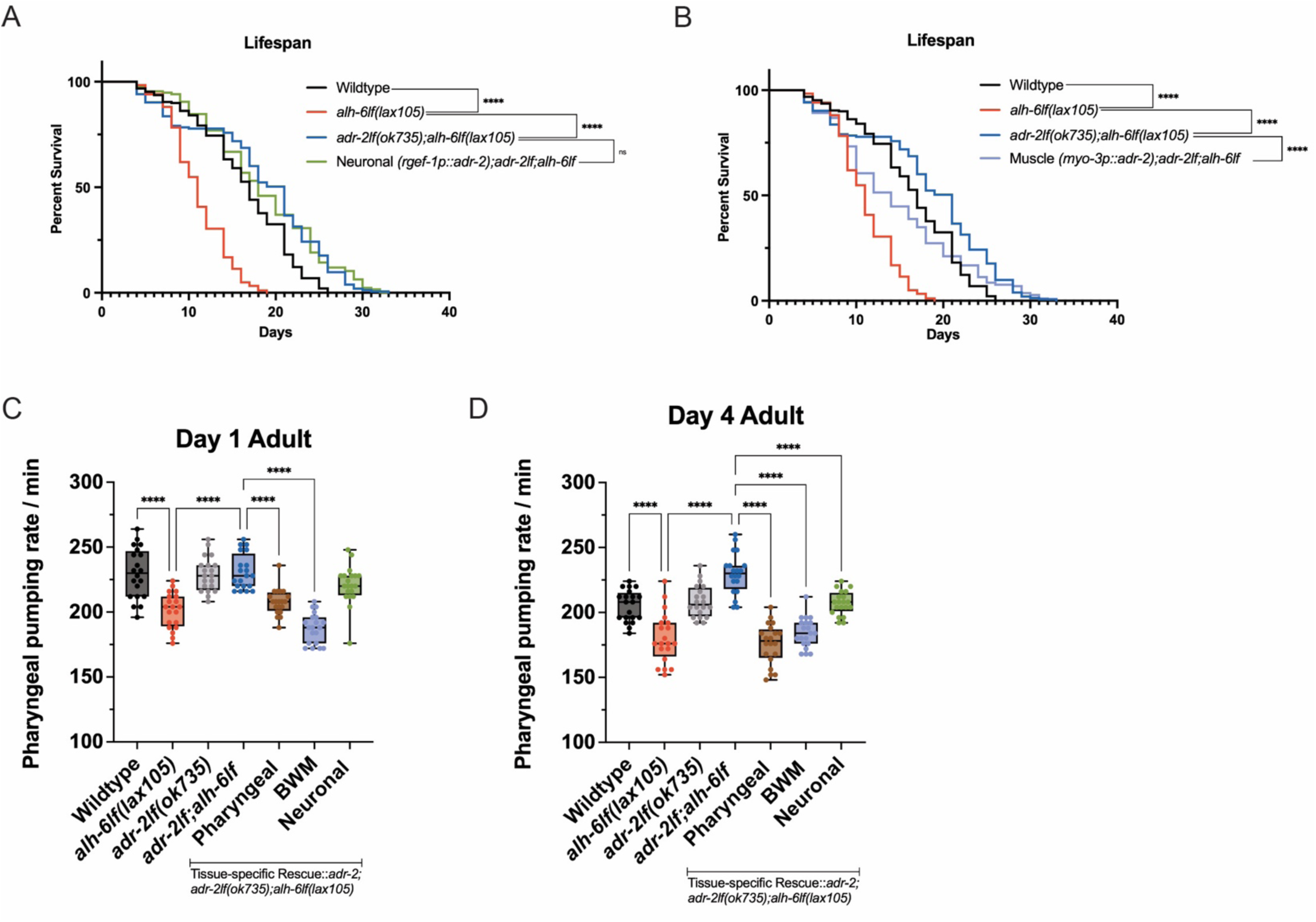
Tissue-specific ADR-2 activity differentially influences lifespan and healthspan in *alh-6lf* mutants. **(A)** Neuronal-specific rescue of *adr-2wt* in *alh-6lf;adr-2lf* animals does not significantly alter lifespan, while **(B)** body wall muscle-specific rescue yields an intermediate lifespan phenotype, with a median survival similar to *alh-6lf* animals but extended survival in the final quartile, resembling *alh-6lf;adr-2lf* mutants. **(C-D)** BWM- and neuronal-specific *adr-2wt* rescues have altered pharyngeal pumping phenotypes; together, these results indicate that ADR-2 mediated RNA editing in the pharynx drive the age-related decline in *alh-6lf* mutants and underlies a cell non-autonomous regulation of healthspan.

**Supplementary Figure S6.**
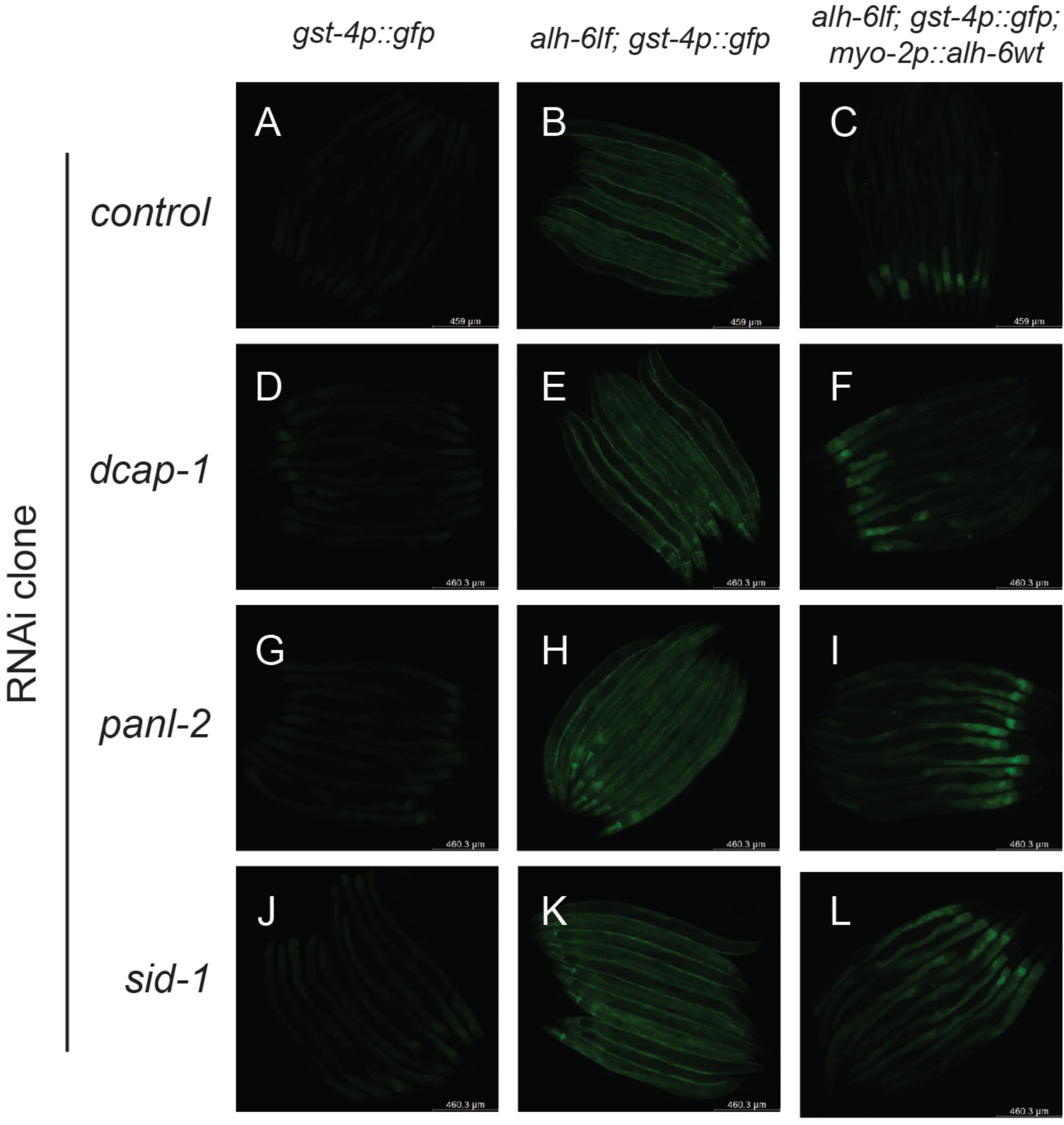
RNA metabolism and transport RNAi genes in *alh-6lf* animals. (A-L) Representative fluorescent images showing *gst-4p::GFP* expression in *alh-6lf* and *myo-2p::alh-6(wt)* animals following RNAi knockdown of *dcap-1, panl-2,* and *sid-1.* Each RNAi induces *gst-4p::GFP* activation outside of the pharynx.

## MATERIALS AND METHODS

### Growth and Maintenance of *Caenorhabditis elegans* Strains

*Caenorhabditis elegans* were raised on 6cm nematode growth media (NGM) plates supplemented with streptomycin and seeded with *E. coli* B strain OP50. All worm strains were grown using standard techniques at 20°C and unstarved for at least three generations before being used.

Strains used in this study include:

N2 Bristol (WT),

Hawaiian CB4856,

CL2166 [*gst-4p::GFP*],

SPC321 [*alh-6(lax105)*],

SPC223 [*alh-6lf(lax105); gst-4p::GFP*],

SPC349 [*adr-2(lax229); alh-6(lax105); gst-4p::GFP*],

BB20 [(*adr-2(ok735)*],

SPC2022 [*(adr-2(ok735); alh-6lf(lax105)*],

SPC2024 [*adr-2(ok735); alh-6lf(lax105);gst-4p::GFP*],

SPC2062 [*drh-1gf(lax257 alh-6(lax105)*],

SPC2063 [*drh-1gf(lax257); alh-6(lax105);gst-4p::GFP*],

SJ4103 [*myo-3p::mitoGFP*],

SPC2064 [*myo-3p::mitoGFP*; *alh-6(lax105)]*,

SPC2056 [SP23[*myo-2p::alh-6::SL2::WrmScarlet::alh-6 3’UTR);alh-6lf(lax105)*]],

SPC2047 [SP25[*rgef-1p::alh-6::SL2::WrmScarlet::alh-6 3’UTR);alh-6lf(lax105*]],

SPC2058 [SP27[*alh-6p::alh-6::SL2::WrmScarlet::alh-6 3’UTR);alh-6lf(lax105*]],

SPC2060 [SP29[*myo-3p::alh-6::SL2::WrmScarlet::alh-6 3’UTR);alh-6lf(lax105)*]],

SPC2048 [SP31[*pie-1p::alh-6::SL2::WrmScarlet::alh-6 3’UTR);alh-6lf(lax105)*]],

SPC2057 [SP23[*myo-2p::alh-6::SL2::WrmScarlet::alh-6 3’UTR);alh-6lf(lax105);gst-4p::GFP*]],

SPC2026 [SP25[*rgef-1p::alh-6::SL2::WrmScarlet::alh-6 3’UTR);alh-6lf(lax105);gst-4p::GFP*]],

SPC2059 [SP27[*alh-6p::alh-6::SL2::WrmScarlet::alh-6 3’UTR);alh-6lf(lax105);gst-4p::GFP*]],

SPC2061 [SP29[*myo-3p::alh-6::SL2::WrmScarlet::alh-6 3’UTR);alh-6lf(lax105);gst-4p::GFP*]],

SPC2054 [SP31[*pie-1p::alh-6::SL2::WrmScarlet::alh-6 3’UTR);alh-6lf(lax105);gst-4p::GFP*]],

SPC625 [SP13 [*myo-2p::adr-2::SL2::WrmScarlet::alh-6 3’UTR);alh-6lf(lax105)*]],

SPC626 [SP15 [*rgef-1p::adr-2::SL2::WrmScarlet::alh-6 3’UTR);alh-6lf(lax105)*]],

SPC630 [SP19 [*myo-3p::adr-2::SL2::WrmScarlet::alh-6 3’UTR);alh-6lf(lax105)*]]

### Oxygen Consumption Rate (OCR) Seahorse Assay

OCR of age-synchronized live animals was measured using an Agilent Seahorse XF96 Analyzer (Agilent Technologies), adapted from Koopman et al., 2016^61^. Briefly, synchronized animals were collected, washed three times with M9 buffer, and allowed to equilibrate for 30 minutes to clear residual bacteria from the gut. Approximately 10-25 animals were transferred into each well of an Agilent islet capture microplate in a final volume of 500mL M9. The exact number of worms per well was recorded and used for normalization of OCR data.

Basal respiration was measured for 5 cycles, followed by injection of the mitochondrial uncoupler FCCP (0.01mM final concentration, from a 5mM DMSO stock) to determine maximal respiration. Subsequently, for non-mitochondrial OCR, sodium azide (40mM final concentration, from a 400mM stock) was injected to inhibit mitochondrial complexes IV and V. OCR was recorded for 2 minutes per measurement cycle, with 9 cycles after FCCP and 4 cycles after sodium azide injection. Raw OCR values were normalized to worm number per well, to yield corrected OCR in pmol O2 min^-1^ worm^-1^. Area under the curve of oxygen consumption rate, before and after the addition of FCCP and sodium azide, was used to determine basal respiration, maximal respiration, and spare respiration capacities, as described in Ng and Gruber, 2019^62^.

### WormLab Measurements

Crawling and thrashing movements were recorded and analyzed as previously described^36^. Briefly, worms were egg prepped with a hypochlorite solution and eggs were allowed to hatch overnight for a synchronous L1 population. The next day, worms were dropped onto plates seeded with OP50. Worms were then allowed to grow until each time point (48 h post-drop for L4s, 72 h post-drop for day 1 adults, and 120 h post-drop for day 3 adults). Once worms were the required stage of development, 30-50 worms were washed off a plate in 50 mL of M9 with a M9 + 0.01% Triton coated P1000 tip and dropped onto an unseeded NGM plate. The M9 was allowed to dissipate, and worms roamed on the unseeded plate for 1 hour before imaging crawling. Crawling was imaged with the WormLab microscope (MBF Bioscience), and analysis was performed with WormLab version 2022. Worm crawling on the plate was imaged for 1 minute for each condition at 7.5 ms. Worm crawling was analyzed with the software and only worms that moved for at least 90% of the time were included in the analysis.

### Lifespan Analysis

Lifespan assays were performed as previously described^8,28,36^. Briefly, worms were egg prepped and eggs were allowed to hatch overnight. The next day synchronized L1 larvae were dropped on NGM plates seeded with OP50. 48 h later, approximately 50 L4 hermaphrodites were moved individually onto new plates in replicates of three for each genotype. Worms were kept at 20°C and transferred every other day during the reproductive period. Worms were scored daily for survival by gentle prodding with a platinum wire. Worms that died of vulva burst, bagging, or crawling to the sides of plates were censored and discarded.

### Microscopy

For imaging of strains, synchronized animals were grown on NGM+strep OP50 seeded plates or RNAi plates seeded with L4440 (empty vector) bacteria or RNAi bacteria, respectively. Animals were imaged at day 1 of adulthood and for aging experiments also imaged at day 3, day 5, day 9, and day 12 of adulthood, or as indicated. For whole-worm imaging: Animals were placed on an agar pad and immobilized with 10 mM sodium azide in M9. Worms in the solution were lined alongside each other, covered with a cover slip, and excess liquid was removed and sealed with nail polish. Worms were imaged with LAS X software and Leica compound microscope equipped with a 63x/1.4 Plan Apochromat objective, GFP filter (11525314), DFC90000 GT camera, and LED5 light source. For RNAi imaging: Animals were placed in a pool of 10 mM sodium azide in M9 on standard NGM plates without bacteria to induce paralysis. Paralyzed animals were lined alongside each other and imaged on a Leica M205FCA automated fluorescent stereomicroscope running LAS X software and equipped with a standard GFP filter, Leica LED3 light source, and Leica K5 camera. For all imaging experiments, 3 biological replicates were performed with 2 technical replicates each, and 1 representative image was chosen for use in figures.

### Mitochondrial Morphology and Imaging

Imaging of muscle mitochondria was performed as described^30^. Worms were immobilized with 10mM sodium azide or 0.10mm Polybead microspheres and mounted onto 2% agarose pads for imaging. Worms in the solution were covered with a cover slip and sealed with nail polish, to prevent evaporation of M9 solution. Worms were imaged with LAS X software and Leica Thunder compound microscope equipped with a 63x/1.4 Plan Apochromat objective, GFP filter (11525314), DFC90000 GT camera, and LED5 light source. Dimensions used are 1024(x) * 1024(y) pixels at a zoom factor of 0.75 and z-step size of 0.50 mm. Mitochondrial morphologies were quantified as described^30,63^. The MitoMAPR macro for FIJI, developed by the William Mair laboratory, was used as this macro has automated methods to quantify and measure various aspects of the mitochondria^33^. To further analyze mitochondrial morphology, shapes of GFP-positive mitochondria were scored as linear/tubular, intermediate, or fragmented as previously described [63].

### EMS Mutagenesis and Insertion-Deletion SNP Mapping

Ethyl methanesulfonate (EMS) mutagenesis was performed as previously described^2,36^. Briefly, SPC223 [*alh-6lf(lax105);gst-4p::GFP*] was mutagenized with EMS for 4 hr at 20°C and dropped on NGM plates seeded with OP50. F2 offspring with reduced GFP expression were selected, indicating suppression of SKN-1(*gst-4*) activation. Isolated mutant was crossed into the polymorphic Hawaiian CB4856 strain, and the strain was isolated and genetically mapped as previously described ^2,8,27,28,64,65^ by using the InDel mapping primer set^38^. Whole-genome sequencing and injection rescue confirmed mutant sequence identity. Mutants were outcrossed to wildtype (N2) to remove background mutations before being used for experiments.

### Whole-genome Sequencing

Whole-genome sequencing (WGS) was performed as previously described^28,36,65^. Briefly, worms were egg prepped and eggs were allowed to hatch overnight. The next day ∼4,000 synchronized L1s were dropped on NGM plates seeded with 25X concentrated OP50. After 48 h, L4 animals were washed three times with M9, homogenized, and genomic DNA was extracted using Zymo Quick-DNA Miniprep Kit (Cat. #D3025). DNA samples were library prepped and sequenced by Novogene. Sequencing data were analyzed using Galaxy workflow: Bowtie2 (Galaxy Version 0.4), Filter (Galaxy Version 0.0.1), MPileup (Galaxy Version 2.0), VarScan (Galaxy Version 2.4.2) and R package DESeq 2.

### Library Construction and Circle-sequencing

Circle-sequencing was carried out on N=4 biological replicates for each genotype and age group. To begin library preparation, 1100 ng of enriched mRNA were fragmented with RNase III (Ambion, AM2290) for 27 minutes at 37°C. RNA fragments were then purified with the Oligo Clean & Concentrator kit (Zymo Research, D4061) according to the manufacturer’s recommendations, except that the columns were washed twice instead of once. The fragmented RNA was then circularized with T4 RNA Ligase 1 in 20 µl reactions (NEB, M0204S) for 2 hours at 25°C, after which the circularized RNA was purified with the Oligo Clean & Concentrator kit (Zymo Research, D4061). The circular RNA templates were then reverse transcribed in a rolling-circle reaction with Superscript III Reverse Transcriptase (Invitrogen, 18080044) by first incubating the RNA for 10 minutes at 25°C to allow the random hexamers used for priming to bind to the templates. Then, the reaction was shifted to 42°C for 20 minutes to allow for primer extension and cDNA synthesis. cDNA templates were purified with the Oligo Clean & Concentrator kit (Zymo Research, D4061) and incubated with the NEBNext Ultra II Non-Directional RNA Second Strand Synthesis kit (NEB, E6111S). The remaining steps for library preparation were then performed with the NEBNext Ultra II DNA Library Prep Kit for Illumina (E7103L) and the NEBNext Multiplex Oligos for Illumina (E6440S) according to the manufacturer’s protocols. Briefly, double-stranded cDNA was entered into end-repair, adapter ligation, and size selected for 500–700 bp inserts using AMPure XP beads (Beckman Coulter, A63880). These molecules were then amplified and barcoded with Q5 DNA polymerase and NEBNext Multiplex Oligos for Illumina (E6440S) using a two-step PCR protocol with a 65°C primer annealing and extension step and a 95°C denaturation step, and using the NEB recommended number of cycles based on samples’ Qubit concentrations. Post-amplification final libraries were cleaned up using AMPure XP beads and sequenced by Novogene on an Illumina Novaseq X Plus to generate 300 base pair paired-end reads.

### Circle-sequencing Analyses

Rolling-circle RNA sequencing reads were processed using a previously developed pipeline for characterizing the SARS-CoV-2 mutational landscape ^66^ and available at https://github.com/jfgout/circseq-seqan2/tree/main. Briefly, reads were first trimmed with fastp v0.24.1 ^67^ using default parameters, except for (1) enabling adapter detection for paired end data, (2) trimming poly-G ends with a minimum size of 6, and (3) only retaining reads with a minimum length of 50 bases after trimming. These reads were then inspected for repeats with at least 20 bases, and each stack of repeats was used to generate a consensus repeat sequence. Consensus repeat sequences were then pseudoaligned to a custom *C. elegans* reference transcriptome with Kallisto v0.52.0^68^. This custom transcriptome consisted of (1) annotated transcripts for the *C. elegans* WS220/ce10 assembly in Ensembl release 66 and (2) annotated transposable elements for ce10 extracted from the UCSC Genome Browser ^69^ Repeatmasker track^70^. After locally optimizing consensus sequence alignments with the seqan2 library^71^, genotypes for each position in each consensus sequence were called, ensuring that (1) the genotype is supported by at least 3 repeats, with all repeats supporting the same base call, (2) the genotype is supported by a cumulative quality score of at least 80, and (3) the first and last 2 bases in each consensus sequence are omitted, since these are more prone to artefactual damage. This analysis yielded one table of high confidence genotype calls at every detected position in the genome, as well as a second superset table of all genotype observations where some of the high confidence genotyping criteria were not met.

Genotypes were further processed in R (v4.3.3). To omit potentially polymorphic sites from the analysis, all genotype observations were inspected and only those at sites with at least 20 reads of support and a reference allele frequency of at least 60% were retained. High confidence genotype calls were then intersected with this filtered list of observations, and only the overlapping high confidence genotype calls were used in downstream analyses. We acknowledge that this filtering step may result in a trade-off between filtering polymorphic sites and filtering *bona fide* ADAR editing sites where the alternative, edited allele may exist at a frequency higher than 40%. The ADAR editing frequency at a site with an ‘A’ reference allele was calculated by dividing the number of ‘G’ calls by the total number of calls at that site. The ADAR editing frequency over a gene was calculated by considering all detected sites with an ‘A’ reference allele within that gene. RNA Polymerase I transcript editing rates were calculated from genes *rrn-1.1*, *rrn-1.2*, *rrn-2.1*, and *rrn-3.1*. RNA Polymerase II transcript editing rates were calculated from nuclear-encoded genes with the protein coding, ncRNA, snoRNA, and snRNA biotype annotations. RNA Polymerase III transcript editing rates were calculated from nuclear-encoded genes with the tRNA biotype annotation. All detected genes of mitochondrial origin were used to calculate mitochondrial RNA polymerase transcript editing rates.

To compare editing frequencies across age and genotype groups, variables were modeled as follows: log10(Mutation Frequency) ∼ Age Group + Genotype Group. Statistical significance was then assessed via rank regression using the Rfit v0.27.0 R package, and a p-value < 0.05 was considered significant. We calculated the enrichment of various genomic features overlapping putative ADAR editing sites using the annotatePeaks script within the Hypergeometric Optimization of Motif EnRichment (HOMER) v5.1 software suite ^72^ using ce10 v7.0 annotations. Functional enrichment analyses of ADAR-edited genes were carried out on the STRING v12.0 platform^73^, using all detectable genes with an ‘A’ allele in either wild-type or *alh-6lf* worms as the background. Pathways and Biological Processes with a false discovery rate (FDR) < 0.05 were considered significant. Gene-level PCA analyses were carried out using the prcomp() function in R, and MDS analyses were carried out by applying the cmdscale(1-spearman correlation) function to the gene editing frequency matrix. To equally weigh each of the genes used in the MDS/PCA plots, we added a pseudocount to all the gene frequencies, defined as 1/largest gene count, and log10 transformed the rates before applying the PCA and MDS functions. To highlight differences in RNA editing, as opposed to differences in gene expression, between groups, PCA and MDS analyses were only carried out on 1175 genes where editing rates could be calculated for all samples and editing was detected in at least 1 sample. We recognize that this approach partially masks group-specific editing patterns, although it allows for a fair assessment of editing rates across groups for a common set of gene targets, and it avoids the analytical issues associated with undefined editing rates due to lack of expression.

### Pharyngeal Pumping Assay

Strains of interest were egg prepped, and eggs were allowed to hatch overnight, the following day synchronized L1 populations were dropped onto NGM+strep OP50-seeded plates. When the synchronized populations reached the desired developmental stage, before imaging, approximately 15-20 worms were singled onto NGM+strep OP50-seeded plates 1 hour to equilibrate before videoing pharyngeal pumping. For pumping analysis, worms were tracked on a microscope for 30 seconds, a video was taken at 10x magnification, tracking the movement of the animal. This procedure was repeated with each of the 20 individual worms, yielding 2 reps of 10 unique videos. This procedure was repeated for Day 1 and Day 4 adult animals. Following the recording of the worms, the videos were slowed and the number of pharyngeal grinder movement over the 30 second video were counted and then multiplied by two in order to calculate the average number of pharyngeal pumps over the span of a minute. Analysis was completed using Excel and GraphPad Prism.

### RNAi Treatment

RNAi treatment was performed as previously described^8,28,36^. Briefly, HT115 bacteria containing specific double-stranded RNA-expression plasmids were inoculated overnight in Luria broth containing selective antibiotic, seeded on NGM plates containing 5mM isopropyl-b-D-thiogalactoside and 50μg ml^-1^ carbenicillin. RNAi was induced at room temperature for 24 h. Synchronized L1 animals were placed on those plates to knockdown indicated genes.

